# Engineered living materials suppress uropathogenic *E. coli* growth and invasion of urothelial cells through sustained probiotic release

**DOI:** 10.64898/2026.09.15.751868

**Authors:** Manivannan Sivaperuman Kalairaj, Veerakit Vanitshavit, Iris George, Leroy Arthur, Mustafa K. Abdelrahman, Mason C. Ansley, Samantha N. Goebel, Konstantinos Stamatis, Mia A. Walling, Isabel K. Murphy, Hayden E. Wylie, Kenneth Hoyt, Philippe E. Zimmern, Sargurunathan Subashchandrabose, Taylor H. Ware

## Abstract

Recurrent urinary tract infection (rUTI) is a significant public health problem. The most common cause of rUTI is uropathogenic *Escherichia coli* (UPEC). Antimicrobial prophylactic and therapeutic regimens for rUTI disrupts the microbiome and leads to infection with antimicrobial resistant organisms. Therefore, there is an urgent need to develop microbiome-sparing alternative approaches for preventing rUTI. Asymptomatic bacteriuria (ABU) *E. coli* strain 83972 (ABEC) outcompetes UPEC in the urinary tract without causing UTI symptoms, but its limited persistence in the bladder restricts its efficacy. Here, we investigate a device made from an engineered living material (ELM) that releases ABEC in a sustained manner as an antibiotic-free platform against UTIs and rUTIs. Using physiologically relevant in vitro models incorporating human urothelial cells, human urine, and periodic urine exchange, we show that ABEC-releasing ELMs suppress UPEC proliferation and inhibit UPEC attachment to and invasion into urothelial cells, particularly when ABEC is maintained at equal or higher levels than UPEC. Because ELMs continuously release ABEC, they sustain competitive pressure even as planktonic bacteria are cleared during voiding, outperforming a single dose of free-floating ABEC. In a rUTI model, sustained ABEC release from ELMs reduces the proliferation and reinvasion of UPEC expelled from infected urothelial cells, while UPEC infiltration into fractured ELMs remains negligible. Finally, we design a first-generation ELM device that can be transurethrally delivered and retained within the mouse urinary bladder, achieving sustained ABEC release in vivo, with ABEC persisting in the bladder, kidneys, and urine for at least 4 days. In summary, we report the development of an ELM device that continuously releases ABEC to suppress UPEC proliferation and urothelial cell invasion in in vitro models of rUTI, and demonstrate sustained ABEC release in vivo in a mouse bladder.

## 1. INTRODUCTION

Urinary tract infections (UTIs) represent one of the most prevalent bacterial infections worldwide, affecting more than 400 million individuals annually [1,2]. In addition to the more common clinical presentation of bladder inflammation, cystitis [3,4], uropathogens can also cause kidney infection or disseminate into the bloodstream, resulting in bacteremia or sepsis [4]. Women are particularly susceptible, with more than 50% experiencing at least one UTI during their lifetime [5]. Importantly, ∼30% of women suffer from recurrent UTI (rUTI) after an initial episode of UTI, imposing substantial morbidity, diminished quality of life, and health care costs. Uropathogenic *Escherichia coli* (UPEC) is the predominant causative agent, responsible for 65–75% of all UTIs [3,6,7]. *Klebsiella* spp., *Staphylococcus* spp., *Enterococcus faecalis, Proteus mirabilis*, and *Candida* spp. are also clinically significant uropathogens [3,6,8–11]. Among UPEC, highly virulent *E. coli* phylogroup B2 strains account for 60–70% of UTIs [12–17], yet some strains within this phylogroup lack type 1 and P fimbriae, and their colonization in urinary tract results in asymptomatic bacteriuria (ABU) rather than symptomatic infection [18–21]. ABU *E. coli* strain 83972 (referred to as ABEC throughout this manuscript for brevity) was originally isolated from a girl with sustained ABU without adverse sequelae and is a naturally avirulent strain lacking key virulence factors [22].

The current standard of care for UTIs is antibiotic therapy [23]. Long-term, low-dose antimicrobial prophylaxis is widely used for effective rUTI prevention. Extended antimicrobial use is increasingly untenable in an era of rising antimicrobial resistance, with prophylaxis-associated resistance rates reaching 80% within 6 months. Allergies and other contraindications against long-term antimicrobial prophylaxis leaves a significant subset of people at high risk for rUTI without any effective management options. Chronic and repeated antimicrobial use is associated with disruption of the host microbiome and further emergence of antibiotic-resistant uropathogens [24–26]. Alternative strategies such as cranberry extract and D-mannose have repeatedly failed to demonstrate significant clinical benefits in high-quality randomized controlled trials [27–30]. Probiotic-based approaches represent a microbiome-sparing alternative for UTI prevention [31].

However, orally administered probiotics have not demonstrated significant reductions in UTI recurrence [31], whereas vaginal or intravesical probiotics have shown more promising results [32–42]. Clinical studies have demonstrated that deliberate inoculation of ABEC reduces the incidence of UTI in patients with neurogenic bladder [34,37]. Consequently, ABEC has emerged as a promising antibiotic-free prophylactic strategy against rUTIs in patients with high residual urine [43]. The efficacy of ABEC is attributed to direct bacterial competition, whereby ABEC effectively outcompetes UPEC and other uropathogens [22,44]. However, the absence of virulence factors in ABEC simultaneously diminishes its ability to persist in the urinary tract of rUTI patients who void normally, limiting its broad adoption as a preventive strategy [43]. Achieving prolonged probiotic persistence at hard-to-access sites such as the urinary bladder remains a significant challenge. We have previously demonstrated that engineered living materials (ELMs) can function as probiotic factories, releasing probiotics in a controlled and sustained manner [45]. Here we evaluate the impact of ABEC-releasing ELMs on controlling uropathogen growth in an in vitro human bladder cell infection model.

ELMs are composites frequently constructed by embedding living microorganisms into organic or inorganic matrices [46–50]. These materials possess complex emergent functionality arising from the interplay between their living and non-living components [46,50,51]. The nonliving component maintains the viability of encapsulated microorganisms by facilitating the diffusion of water, nutrients, gases, and biomolecules [46]. Notably, the properties of the nonliving matrix modulate the interactions of the microbes with the surrounding environment [49,52]. Encapsulated microorganisms can utilize the nutrients that diffuse into the matrix to proliferate within ELMs [45,48,52,53]. In many ELMs, this microorganism growth has been shown to result in microbial escape from the matrix [45,48,51,54,55]. We have previously demonstrated the mechanism through which microorganisms, including ABEC, escape from ELMs [45]. Moreover, we showed that this mechanism enables sustained release of microorganisms, approaching zero-order release kinetics, and that the release can be precisely controlled [45]. This precise control over microorganism release could be exploited to realize clinical utility. Specifically, by releasing clinically relevant doses of probiotics, it may be possible to achieve probiotic persistence at hard-to-access sites such as the bladder, where probiotics do not generally colonize but where their persistence could offer significant health benefits.

To test the ability of probiotic-releasing ELMs against UTIs, the material must be engineered for delivery and retention within the bladder. The non-living matrices of ELMs are typically composed of hydrogels, a class of biomaterials extensively employed in implantable devices owing to their potential for biocompatibility. Implantable ELMs have been explored for drug delivery [56]. To date, however, no ELM-based bladder-resident implant has been reported. Other bladder-resident devices are used for applications including urodynamic sensing and sustained intravesical drug delivery for bladder cancer [57]. These devices employ specific design strategies to achieve long-term retention within the bladder. One such strategy involves shape-morphing implants that are straight during catheter-assisted insertion through the urethra but transform into three-dimensional geometries once deployed within the bladder [57]. These expanded forms exceed the dimensions of the bladder neck, preventing expulsion during voiding, while incorporating open regions that permit unobstructed urine flow.

Here, we investigate the potential of ELMs as a prophylactic platform against UTIs and rUTIs. Using in vitro models that incorporate human urothelial cells, urine and periodic urine exchange to mimic voiding, we evaluate the capacity of ABEC-releasing ELMs to suppress UPEC growth in the urine and to limit UPEC adhesion to and invasion into urothelial cells. Although in vitro studies have evaluated UTI prevention strategies using urothelial cells, they were performed in cell culture media rather than in human urine [58,59], in which ABEC fails to outcompete UPEC [43]. Separately, we design shape-changing ELMs suitable for transurethral delivery via catheter and retention within the bladder in a mouse model and assess their ability to release ABEC. Together, these studies establish a framework for achieving sustained probiotic persistence in the bladder and set the stage for further evaluation of ELM-based devices as a precision microbial modulation strategy for the prevention of rUTIs (**Figure 1A**).

**Figure 1.**
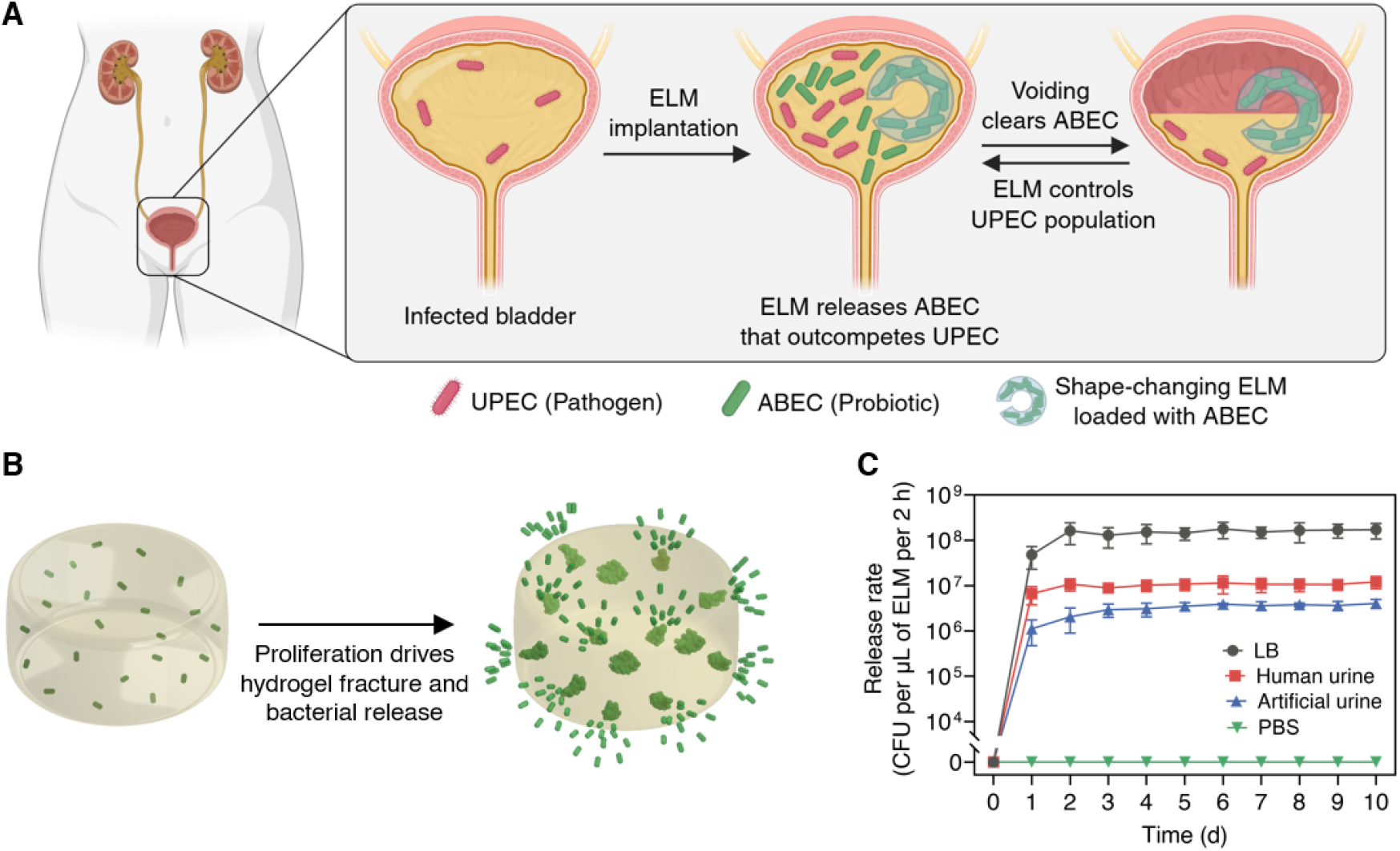
Sustained release of probiotics from ELMs for treating UTIs. **(A)** Schematic illustrating that sustained release of ABEC from shape-changing ELMs could be used to restrict UPEC proliferation in the bladder. **(B)** Schematic illustrating the mechanism of sustained release of ABEC from ELMs by inducing fractures in the hydrogel. **(C)** ABEC release rate as a function of time from ELMs in different milieu. ELMs were prepared with ∼1 × 10^5^ CFU of ABEC per μL of ELM and incubated in 2 mL of the indicated media at 37 °C with shaking at 200 rpm under aerobic conditions. Media was refreshed every 24 h. Data are presented as mean ± standard error of the mean (*n* = 3). Trend lines are only intended to guide the eye.

## 2. RESULTS AND DISCUSSION

### 2.1. ELMs release probiotics in a sustained manner

ELMs can be used to fabricate probiotic factories that release probiotics in a controlled and sustained manner over a prolonged period. To create a sustained release device, ELMs were synthesized by encapsulating living probiotics (ABEC) within acrylic hydrogels. The hydrogels were prepared by free radical polymerization of 2-hydroxyethyl acrylate (HEA) (monomer) and *N,N*′-methylenebis(acrylamide) (BIS) (cross-linker). We have previously demonstrated that these ELMs release ABEC in a sustained manner when incubated in Luria-Bertani (LB) media at 37 °C under aerobic conditions [45]. In this system, the relatively high elastic modulus bacteria proliferate within the relatively low elastic modulus hydrogel matrix in the presence of nutrients, inducing fractures that facilitate probiotic release (**Figure 1B**). During the first 2–3 days, the number of fractures increases, leading to an increase in the rate of release. Subsequently, a steady state is reached where no new fractures are likely occurring, and the number of cells being released is proportional to the number of cells present within the ELMs, primarily controlled by available nutrients and the doubling time of the microorganisms [45]. A similar steady-state release trend is observed with ELMs prepared with 1 × 10^5^ CFU of ABEC per μL of ELM (**Figure 1C**). When these ELMs were incubated in human urine and synthetic urine, a similar near-constant release profile is observed. The probiotic release in human and synthetic urine is more than an order of magnitude lower compared to LB media. This difference in probiotic release can be attributed to the lower nutrient availability and increased osmotic stress in human and synthetic urine, which increases the doubling time of ABEC [43].

### 2.2. UPEC do not colonize ELMs effectively

The fractures in ELMs do not facilitate substantial UPEC infiltration. We have previously demonstrated that ELMs release probiotics in a sustained manner through a fracture-based mechanism [43,45,52]. However, those studies were conducted in sterile growth media in the absence of pathogens. We evaluate whether pathogens can attach to or infiltrate the ELMs (**Figure 2A**), inadvertently increasing pathogen colonization. To assess potential UPEC colonization of the ELM, cell-free hydrogels and 3-day-grown ELMs (living and dead) were incubated in human urine supplemented with ∼1 × 10^6^ CFU/mL of UPEC. After this co-culture, the ELM was homogenized, and bacterial load was enumerated. We note that this method cannot distinguish between infiltrated and attached UPEC. Within 24 h, UPEC counts in the urine increase to > 1 × 10^8^ CFU/mL across all conditions (**Figure S1A**). Somewhat higher UPEC counts are observed in both living (8.89 ± 2.09 × 10^3^ CFU per μL of ELM) and dead ELMs (8.45 ± 2.31 × 10^3^ CFU per μL of ELM) compared to cell-free hydrogels (1.88 ± 0.68 × 10^3^ CFU per μL of ELM, *P* < 0.05) (**Figure 2B**). There is no significant difference in UPEC colonization between living and dead ELMs.

**Figure 2.**
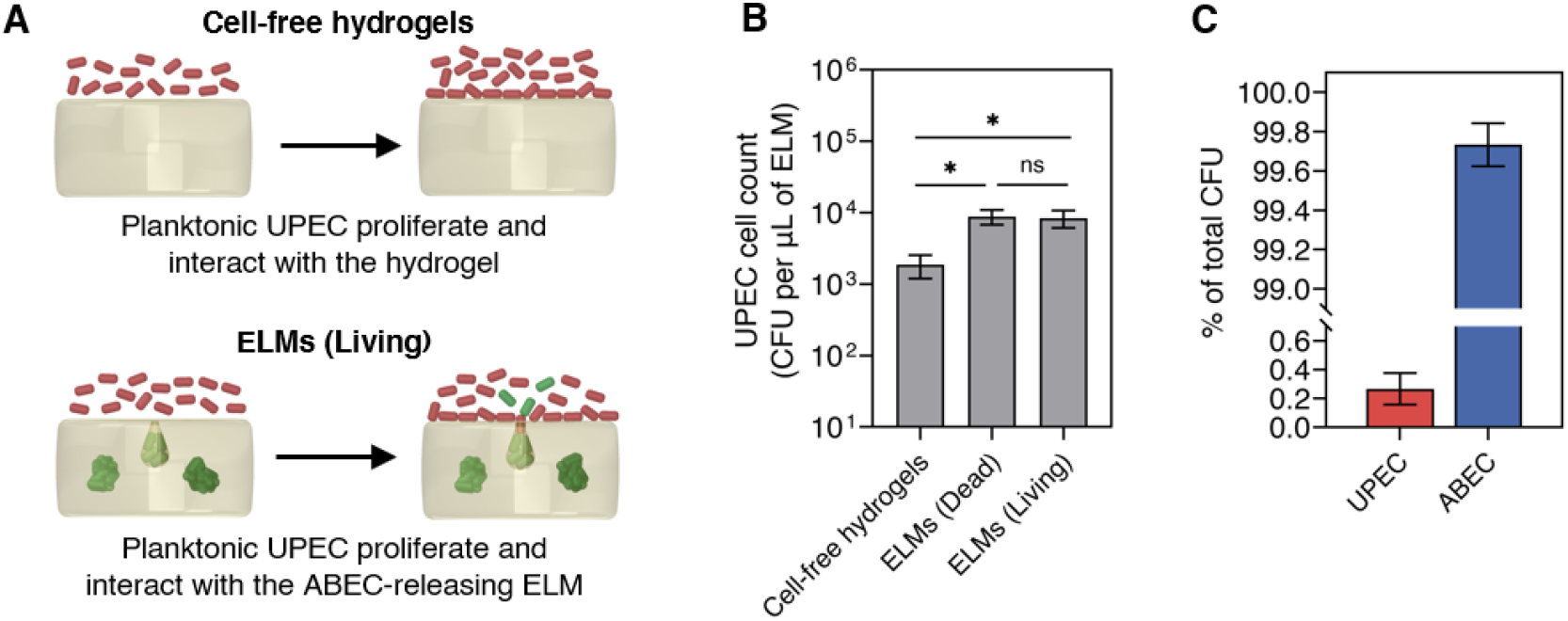
UPEC interaction with ELMs. **(A)** Schematic illustrating potential UPEC association with cell-free hydrogels or ABEC-releasing ELMs. **(B)** UPEC associated with cell-free hydrogels and ELMs (attached to or within) after 24 h of incubation with UPEC. **(C)** Relative abundance of ABEC and UPEC present in ELMs (living) after 24 h of incubation with UPEC. Cell-free hydrogels and 3-day-grown ELMs (both living and dead) were incubated in 2 mL of human urine supplemented with ∼1 × 10^6^ CFU/mL of UPEC at 37 °C with shaking at 200 rpm under aerobic conditions for 24 h. Data are presented as mean ± standard error of the mean (*n* = 5). Statistical analysis in panel **(B)** was performed using a Kruskal–Wallis test with Dunn’s multiple comparisons test. * *P* ≤ 0.05 and not significant (ns) for *P* > 0.05.

UPEC colonization of living ELMs is minimal (0.27 ± 0.11%) compared to the ABEC population (99.73 ± 0.11%) within ELMs (**Figure 2C, Figure S1B**). Since cell-free hydrogels do not contain fractures or pores similar in size to UPEC, UPEC may attach to their surface, but they are not expected to infiltrate the gel. Studies have demonstrated that bacterial migration and growth in confined spaces are limited [60], and microbial infiltration is minimal without designed porosity [61]. Living and dead ELMs have fractures that could serve as infiltration sites. Living ELMs release ABEC through those fractures in a sustained manner (**Figure S1C**); dead ELMs do not release ABEC but still retain fractures that could permit infiltration. The lack of difference between living and dead ELMs with regard to UPEC infiltration or attachment potentially suggests that infiltration through fractures is not the primary mechanism observed. The moderately increased UPEC association with ELMs compared to cell-free hydrogels could also be attributed to the larger surface area of grown ELMs relative to cell-free hydrogels [45,48,52]. In any case, the limited colonization of UPEC into ELM fractures suggest that pathogen association with ELM is unlikely to affect probiotic release from ELM, enabling future development of ELMs as intravesical devices.

### 2.3. ABEC-ELMs suppress UPEC proliferation in urine

The presence of ABEC-releasing ELMs in urine prior to UPEC infection effectively reduces the pathogen population within 6 h of competition in the presence of urothelial cells (**Figure 3A**). ELMs were first cultured for 3 days in human urine to ensure a steady release of ABEC. When 3-day-grown ELMs were placed in urine with uninfected 5637 human urothelial monolayer, ELM-released ABEC population reached 1.31 ± 0.44 × 10^6^ CFU/mL of urine in the first 2 h, increasing to 1.91 ± 0.32 × 10^8^ CFU/mL over the next 6 h, even when urine was refreshed every 2 h (**Figure S2**). UPEC strain CFT073 is used as a prototypical clinical strain for investigating host-uropathogen interaction. However, its hemolysin rapidly damages and kills urothelial cells [62,63], precluding competition experiments in the presence of urothelial cells beyond 2 hours. We used a hemolysin-deficient mutant (CFT073 Δ*hly*::Cam) to overcome this limitation. To evaluate the prophylactic efficacy, we tested: (1) ABEC-releasing ELMs co-cultured with UPEC, (2) free-floating ABEC co-cultured with UPEC, and (3) a control containing only UPEC (without ABEC or ELMs). For the ELM condition, 3-day-grown ELMs that release 1.25 ± 0.21 × 10^6^ CFU/mL in 2 h were introduced at the –2 h timepoint. For the free-floating condition, an equivalent concentration of ABEC (1.25 ± 0.21 × 10^6^ CFU/mL) was introduced at the 0 h timepoint. In all three conditions, different concentrations of UPEC were introduced at 0 h, and urine was refreshed every 2 h for 6 h (**Figure 2B, Figure S3**). Four ABEC-to-UPEC ratios of 100:1, 10:1, 1:1, and 1:10 were examined.

**Figure 3.**
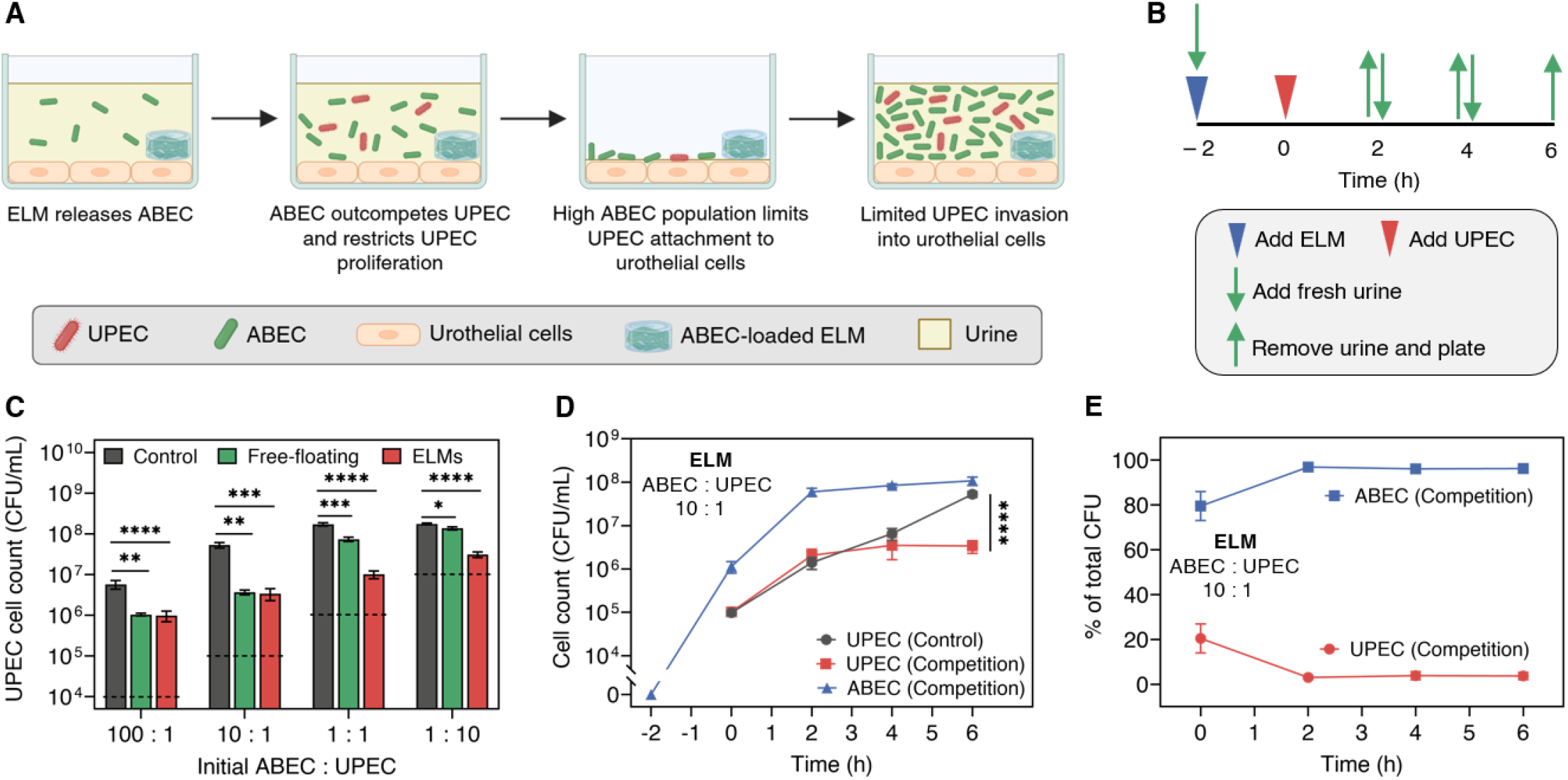
Prophylactic ELMs suppress UPEC proliferation. **(A)** Schematic illustrating that sustained release of ABEC from ELMs restricts UPEC proliferation in urine in the presence of urothelial cells. **(B)** Timeline of the competition experiments. **(C)** UPEC count in the supernatant (urine) after 6 h of competition in free-floating and ELM conditions compared to controls at different initial ABEC:UPEC ratios (100:1, 10:1, 1:1, and 1:10). **(D)** ABEC and UPEC colony-forming unit (CFU) counts in the supernatant (urine) over 6 h of competition in ELM conditions compared to controls at an initial ABEC:UPEC ratio of 10:1. **(E)** Relative abundance of ABEC and UPEC in the supernatant (urine) over 6 h of competition in the ELM condition at an initial ABEC:UPEC ratio of 10:1. The dashed line in panel **(C)** represents the initial UPEC concentration added to the respective ratios in all conditions (control, free-floating, and ELM). Experiments were performed in 24-well plates incubated at 37 °C with 5% CO_2_. ELMs were added at –2 h, UPEC and free-floating ABEC were introduced at 0 h, and human urine was refreshed every 2 h for 6 h. Data are presented as mean ± standard error of the mean (*n* = 9). Statistical analysis in panels **(C)** and **(D)** was performed using a two-tailed Mann–Whitney U test. * *P* ≤ 0.05, ** *P* ≤ 0.01, *** *P* ≤ 0.001, **** *P* ≤ 0.0001, and not significant (ns) for *P* > 0.05. Trend lines are only intended to guide the eye.

ELM-released ABEC to UPEC at 100:1 and 10:1 ratios significantly suppress UPEC proliferation. UPEC proliferation over 6 h in our in vitro infection model is significantly higher in the control condition than in both the free-floating and ABEC-ELM conditions (**Figure 3C**,**D, Figure S4, Figure S5**). This reduced UPEC proliferation in the presence of ABEC can be attributed to the competition between UPEC and ABEC. During this co-culture, ABEC population increases in both the free-floating and ELM conditions (**Figure 3D, Figure S4, Figure S5**). Overall, both conditions effectively suppress expansion of UPEC populations within 6 h. In the free-floating and ELM conditions at 100:1 ratio, the UPEC fraction remains near 1% **(Figure S4C, Figure 3E)**. In the free-floating and ELM condition at 10:1 ratio, the UPEC fraction decreases to around 3%. (**Figure S4D, Figure S5B**).

Interestingly, ELMs releasing equal or fewer ABEC compared to the UPEC suppress UPEC proliferation better than free-floating ABEC. At the 1:1 and 1:10 ratios, UPEC proliferation over 6 h is significantly higher in the control than in both the ELM and free-floating conditions (**Figure 3A**). During this co-culture, ABEC population increases in both the free-floating and ELM conditions (**Figure S6, Figure S7**). Although both conditions reduce the UPEC population compared to controls, the magnitude of suppression differs dramatically. In the free-floating condition, the UPEC fraction decreases only from 47 ± 3% to 41 ± 3% at the 1:1 ratio and from 89 ± 1% to 78 ± 2% at the 1:10 ratio over 6 h. The ABEC-ELM achieves substantially greater reductions, from 56 ± 11% to 9 ± 1% and from 85 ± 6% to 32 ± 4% at the 1:1 and 1:10 ratios, respectively (**Figure S6, Figure S7**). This difference could be attributed to the higher ABEC load from continuous release from the ELM, which maintains ABEC levels an order of magnitude higher than the free-floating ABEC. The lower numbers of free-floating ABEC are unable to outcompete the high UPEC population, particularly at the 1:10 ratio. This aligns with our previous results showing that free-floating ABEC fail to outcompete UPEC when present at a substantial population disadvantage [43]. In contrast, the continuous release of ABEC from ELMs sustains competition for a longer period, eventually emerging as the dominant strain in the total population.

The increased ability of ABEC-ELMs compared to free-floating ABEC to reduce UPEC proliferation can be attributed to the sustained high loads of the ABEC within the co-culture. Since ABEC do not adhere effectively to the urothelial cells [19], urine refresh cycles resulted in greater loss of free-floating ABEC, diminishing its ability to outcompete the higher UPEC numbers. By contrast, continuous release from ELMs maintains competitive pressure against UPEC, exerting substantially greater control over expansion of UPEC population. The suppression of UPEC is driven by sustained ABEC release rather than the hydrogel matrix itself. When cell-free hydrogels were tested under the same conditions, no significant difference in UPEC population is observed between the cell-free hydrogel and control without ABEC in culture (**Figure S8**). We note that the urine was refreshed manually using a pipette, which does not accurately simulate physiological urine flow. Under more physiologically accurate flow conditions, we expect even further ABEC loss in free-floating conditions, decreasing its ability to control UPEC population.

### 2.4. ABEC-ELMs decrease UPEC adherence and invasion in urothelial cells

UPEC adherence and invasion in a human urothelial cell line were determined after 6 h of competition with ABEC-ELMs or controls (**Figure 4A**). When ELMs release equal or higher numbers of ABEC compared to the UPEC (100:1, 10:1, and 1:1 ratios), UPEC attachment to the urothelial cells is significantly lower with ELMs compared to the controls (**Figure 4B**). This reduction in UPEC attachment can be attributed to the high ABEC population in the supernatant and higher numbers of ABEC associated with urothelial cells observed in the ELM condition (**Figure S9A**). The high viable ABEC counts in the supernatant, one to two orders of magnitude greater than UPEC across all ratios, continuously outcompete UPEC. Since the experiment was performed under static conditions, these abundant ABEC can settle onto the urothelial cells decreasing UPEC access to receptors for adhesion. Although ABEC lack well characterized adhesins including the type 1 and P fimbriae, urothelial cell-associated ABEC count exceeds that of UPEC by more than two orders of magnitude at the 100:1 ratio, more than one order of magnitude at the 10:1 ratio, and approximately half an order of magnitude at the 1:1 ratio, likely due to non-specific interaction. When ELMs release fewer ABEC compared to the UPEC population (1:10 ratio), the effectiveness of ELMs in reducing UPEC attachment diminishes (**Figure 4B**). No significant difference in UPEC attachment is observed between the control and ELM conditions. The free-floating ABEC also significantly reduces UPEC attachment at the 100:1, 10:1, and 1:1 ratios compared to the control, with no significant difference observed at the 1:10 ratio (**Figure S9B**). Together, these results indicate that maintaining a high ABEC load in the supernatant can dramatically reduce UPEC attachment to urothelial cells.

**Figure 4.**
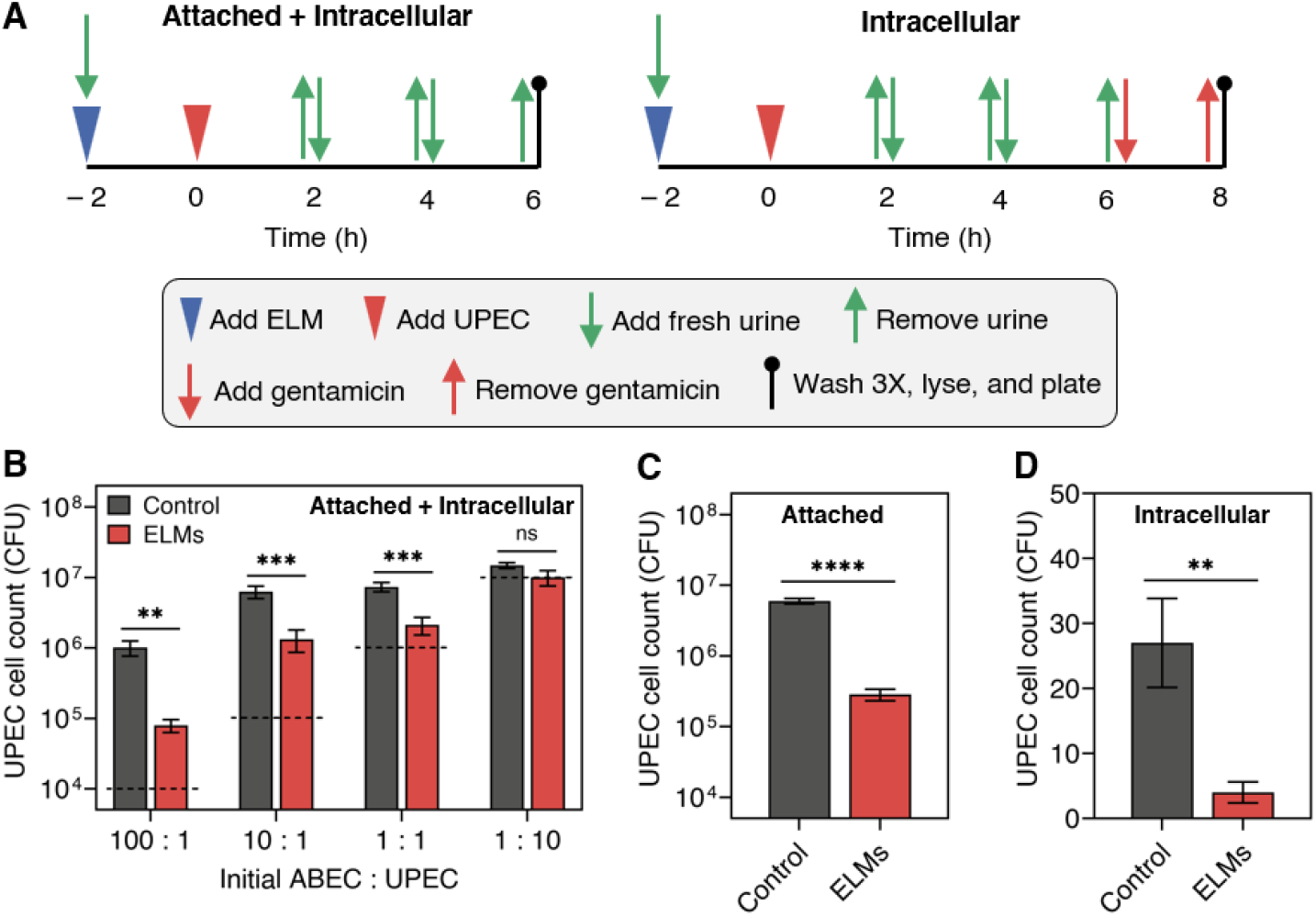
Prophylactic ELMs inhibit UPEC attachment to and invasion into urothelial cells. **(A)** Timeline of the competition experiment performed in 24-well plates in the presence of urothelial cells for ELM condition, depicting the quantification of attached and invaded bacterial cells. **(B)** UPEC counts in the lysate (including both attached and intracellular UPEC) after 6 h of competition in ELM conditions compared to control conditions at different initial ABEC:UPEC ratios (100:1, 10:1, 1:1, and 1:10). **(C)** UPEC counts attached to the urothelial cells in control and ELM conditions after 6 h of competition at an initial ABEC:UPEC ratio of 10:1. **(D)** UPEC counts invaded into urothelial cells in control and ELM condition after 6 h of competition at an initial ABEC:UPEC ratio of 10:1. The dashed line in panel **(B)** represents the initial UPEC concentration added to the respective ratios in both conditions (control and ELM). Experiments were performed in 24-well plates incubated at 37 °C with 5% CO_2_. ELMs were added at –2 h, UPEC were introduced at 0 h, and human urine was refreshed every 2 h for 6 h. Data are presented as mean ± standard error of the mean (*n* = 9). Statistical analysis in panels **(B)–(D)** was performed using a two-tailed Mann–Whitney U test. * *P* ≤ 0.05, ** *P* ≤ 0.01, *** *P* ≤ 0.001, **** *P* ≤ 0.0001, and not significant (ns) for *P* > 0.05.

A high concentration of extracellular ABEC inhibits UPEC invasion into urothelial cells. To quantify invasion after 6 h of ABEC-UPEC competition, cells were treated with gentamicin for 2 h to eliminate extracellular bacteria and intracellular bacterial load was determined (**Figure 4A**). Adherent counts were determined by subtracting the intracellular CFU from the total lysate CFU yield. At the 10:1 ratio, UPEC attachment (**Figure 4C**) and invasion (**Figure 4D**) to urothelial cells are significantly lower with the ABEC-ELMs than controls. This marked reduction in UPEC attachment and invasion can be attributed to the lower UPEC population in the supernatant and the high number of ABEC competing for nutrients and possibly preventing access to receptors. Although ABEC do not invade urothelial cells, the high ABEC association with urothelial cells (**Figure S9A**) and the high ABEC presence in the urine deters UPEC attachment. Since UPEC attachment is a prerequisite for invasion [64], the ABEC-mediated reduction in attachment directly translates to reduced UPEC invasion. Since UPEC invasion into urothelial cells is the primary driver of rUTIs [65], ABEC-releasing ELMs might be a promising strategy for controlling rUTIs.

### 2.5. ABEC inhibits proliferation of UPEC expelled from intracellular reservoirs

The presence of ABEC in urine outcompetes expelled UPEC, reducing extracellular UPEC proliferation (**Figure 5A**). UPEC CFT073 and UTI89 are widely used prototypical clinical strains [66]. UTI89 is well characterized for its ability to invade urothelial cells and form intracellular reservoirs [67] without causing excessive cytotoxicity. Therefore, we used UTI89 to form intracellular reservoirs to test the effect of ABEC-ELMs on preexisting UPEC reservoirs. To initiate UPEC invasion, UTI89 (at a multiplicity of infection (MOI) of 100) was added to urothelial cells and incubated for 30 min, followed by washing to remove non-adherent bacteria (**Figure 5B**). Gentamicin was subsequently added for 2 h to eliminate any non-invaded UPEC. After invasion, urothelial cells were lysed to measure the initial intracellular UPEC count (795 ± 121 CFU). Urine was then added to the urothelial cells. Urothelial cells are capable of rapidly expelling internalized UPEC back into the extracellular environment [58,68,69]. These expelled UPEC proliferate in the urine, enabling them to re-attach to and re-invade urothelial cells (**Figure S10A**).

**Figure 5.**
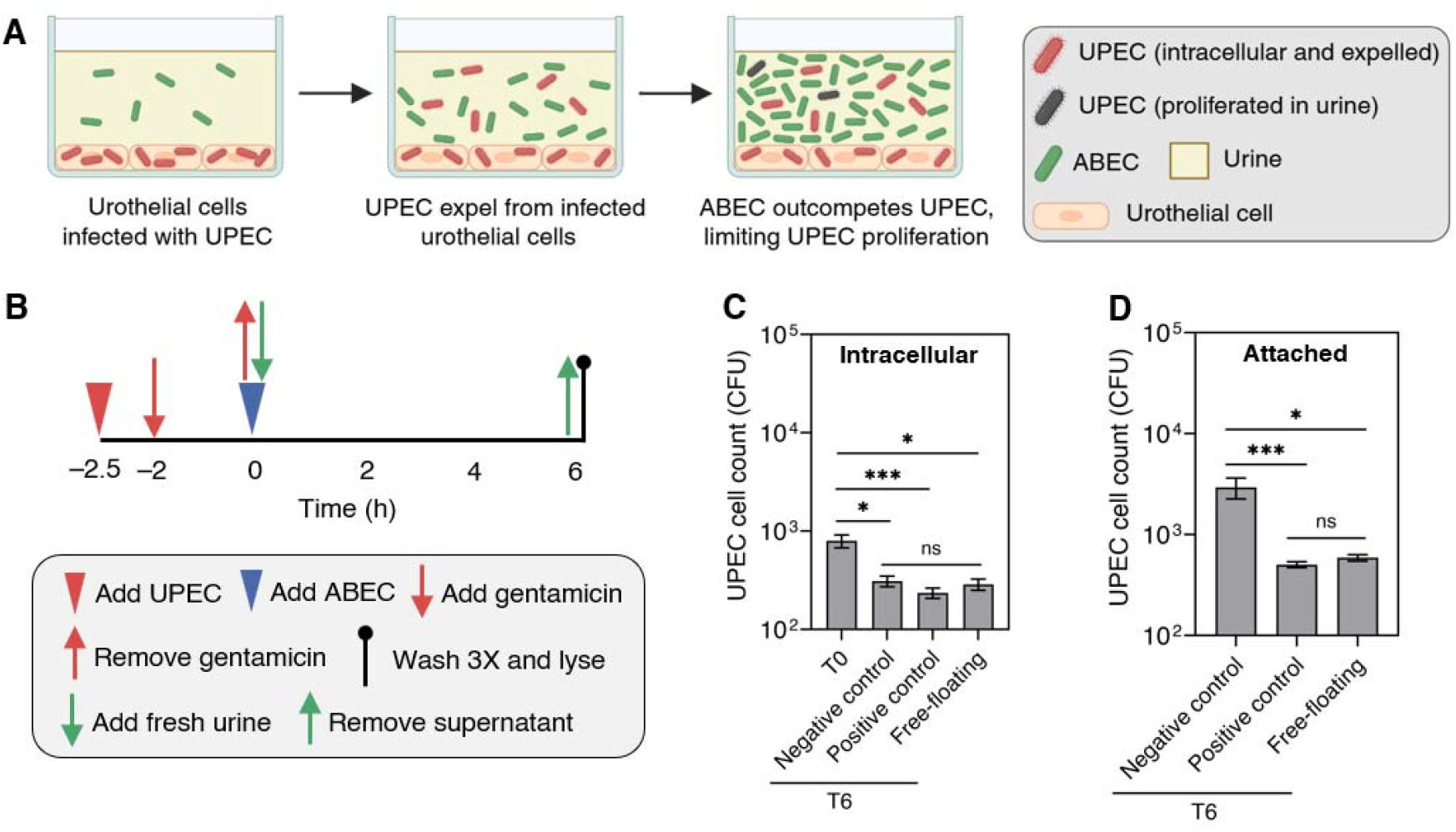
ABEC inhibits proliferation of expelled UPEC. **(A)** Schematic illustrating that ABEC present in supernatant restricts proliferation of expelled UPEC and reduces attachment to urothelial cells. **(B)** Timeline of the invasion experiment performed in 24-well plates in the presence of urothelial cells for the free-floating ABEC condition. **(c)** ABEC and UPEC counts in the supernatant (urine) over 6 h of invasion experiment in free floating conditions compared to control conditions. **(d)** ABEC and UPEC counts in the lysate (including both attached and intracellular UPEC) over 6 h of invasion experiment in free-floating conditions compared to control conditions. **(C)** UPEC counts present intracellularly in the urothelial cells at 0 h and 6 h for three different conditions (negative control, positive control, and free-floating). **(D)** UPEC counts attached to the urothelial cells after 6 h of invasion experiment in three different conditions (negative control, positive control, and free-floating). Experiments were performed in 24-well plates incubated at 37 °C with 5% CO_2_. Free-floating ABEC (free-floating condition) or trimethoprim with methyl α-D-mannoside (positive control condition) were added post-invasion. Human urine was not refreshed for 6 h in all conditions. Data are presented as mean ± standard error of the mean (*n* = 9). Statistical analysis in panels **(C)** and **(D)** was performed using a Kruskal–Wallis test with Dunn’s multiple comparisons test. * *P* ≤ 0.05, ** *P* ≤ 0.01, *** *P* ≤ 0.001, **** *P* ≤ 0.0001, and not significant (ns) for *P* > 0.05.

We evaluate the efficacy of ABEC against intracellular UPEC using (1) free-floating ABEC (1.07 ± 0.05 × 10^6^ CFU/mL), (2) a positive control with a bacteriostatic antibiotic (trimethoprim) to prevent UPEC growth and methyl α-D-mannoside to prevent reattachment of the expelled UPEC to the urothelial cells [59], and (3) a negative control containing only infected urothelial cells (**Figure S10B**). Within 6 h of the expulsion assay, the intracellular UPEC decrease significantly in all conditions (**Figure 5C**), indicating expulsion of intracellular UPEC. Moreover, no significant difference in the intracellular UPEC count after 6 h is observed among the three conditions, suggesting that UPEC expulsion is similar regardless of condition and is likely not driven by the presence of ABEC or bacteriostatic antibiotics.

After 6 h, UPEC proliferation in the urine in the negative control is significantly higher than in both the positive control and free-floating ABEC (**Figure S10C**). In the positive control, expelled UPEC are unable to proliferate due to the bacteriostatic antibiotic. In the free-floating condition, ABEC population increases within 6 h, effectively outcompeting and reducing UPEC proliferation (**Figure S10D**). Extracellular UPEC in the supernatant can attach to urothelial cells [59]. Within 6 h, UPEC attachment in the negative control condition is significantly higher than in both the positive control and free-floating conditions (**Figure 5D**). No significant difference is observed between positive control and free-floating conditions. The reduced UPEC attachment in the positive control can be attributed to the presence of methyl α-D-mannoside. In the free-floating condition, the high ABEC in the supernatant and its association with the urothelial cells decreases UPEC attachment (**Figure S10D, Figure S11**).

ABEC reduces both attached and intracellular UPEC levels comparable to those achieved by a bacteriostatic antibiotic. Since intracellular UPEC is the primary contributor of rUTIs [65,70], these results suggest that maintaining a high ABEC population in the supernatant is a potential strategy to reduce rUTIs. We note that the high ABEC load in the supernatant can be attributed to the absence of urine refreshment over the 6 h experiment. ABEC is cleared from the bladder over time after a single administration of free-floating ABEC since it does not establish persistence in the bladder [43]. This motivates the development of implantable ELMs that release ABEC in a sustained manner to address this gap.

### 2.6. ELMs inhibit proliferation of expelled UPEC

ABEC released from ELMs outcompete expelled UPEC, reducing extracellular UPEC proliferation (**Figure 6A**). Following the invasion procedure described in Section 2.5, UTI89 was used to establish intracellular UPEC colonization in urothelial cells, resulting in an intracellular UPEC count of 2.44 ± 0.24 × 10^3^ CFU. Two conditions were tested: (1) a control for expelled UPEC from intracellular reservoir (**Figure S12A**), and (2) an ABEC-ELM condition. In both conditions, urine was refreshed every 2 h, and the experiment was performed for 6 h (**Figure 6B, Figure S12B**). After 6 h, expelled UPEC proliferates to higher levels in urine of controls than the ABEC-ELM group (*P* < 0.0001) (**Figure 6C**). These extracellular UPEC can re-attach to and re-invade urothelial cells [71]. UPEC counts in the lysate (including attached and intracellular UPEC) are significantly higher in the controls than in the ABEC-ELM at 6 h (*P* < 0.05) (**Figure 6D**), corresponding to a 55 ± 10% UPEC reduction with ABEC-ELMs (**Figure 6E**). This reduction is caused by the high ABEC levels in the supernatant and association with urothelial cells (**Figure S12C**). No significant difference in UPEC load is observed between the cell-free hydrogel and control conditions in either the supernatant or lysate (**Figure S13**). These results indicate that sustained release of ABEC from ELMs is an effective strategy to slow proliferation of expelled UPEC from intracellular reservoirs. Our findings set the stage for future studies to evaluate whether deterrence of growth, re-adherence, and re-invasion of UPEC by ABEC-ELMs could eliminate intracellular reservoirs over time in an animal model of rUTI.

**Figure 6.**
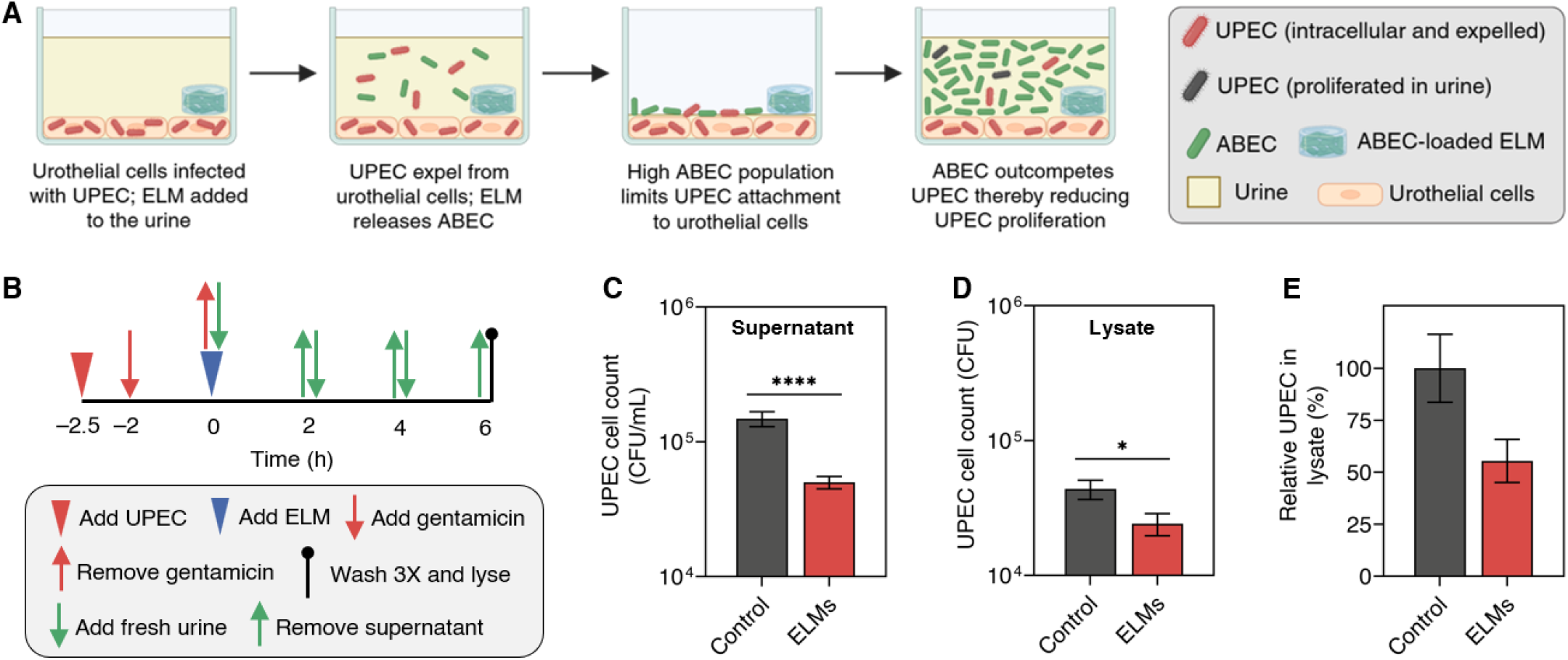
ELMs inhibit proliferation of expelled UPEC. **(A)** Schematic illustrating that sustained release of ABEC from ELMs restricts proliferation of expelled UPEC and reduces their attachment to urothelial cells. **(B)** Timeline of the invasion experiment performed in 24-well plates in the presence of urothelial cells for the ELM condition. **(C)** UPEC counts in the supernatant (urine) after 6 h of the invasion experiment in the ELM condition compared to the control condition. **(D)** UPEC counts in the lysate (including both attached and intracellular UPEC) after 6 h of the invasion experiment in the ELM condition compared to the control condition. **(E)** Relative UPEC counts in the lysate after 6 h of the invasion experiment in the ELM condition compared to the control condition. Experiments were performed in 24-well plates incubated at 37 °C with 5% CO_2_. ELMs were added post-invasion, and human urine was refreshed every 2 h for 6 h. Data are presented as mean ± standard error of the mean (*n* = 10). Statistical analysis in panels **(C)** and **(D)** was performed using a two-tailed Mann–Whitney U test. * *P* ≤ 0.05, **** *P* ≤ 0.0001, and not significant (ns) for *P* > 0.05.

### 2.7. Shape changing ELMs release probiotics in a sustained manner

Since our future goal is to test ABEC-ELMs for preventing rUTIs, we wanted to develop ABEC-ELM devices that could be safely delivered to the bladder, retained without obstruction, and naturally eliminated from the bladder over time. Bladder-resident devices have been developed using design strategies that achieve long-term retention [57]. One such strategy involves shape-morphing implants that are straight during catheter-assisted insertion through the urethra and transform into three-dimensional geometries once deployed in the bladder [57]. The expanded forms exceed the dimensions of the bladder neck to prevent expulsion during voiding, while incorporating open regions that permit unobstructed urine flow.

Shape-morphing devices can be developed using hydrogel-based bilayers or trilayers, in which differential swelling between layers drives bending [72,73] (**Figure 7A**). We designed a trilayer intravesical ELM-device in which a hydrogel bilayer (175 μm thick) was prepared with 69 HEA/2 AM/2 BIS/25 CNC. The CNCs were allowed to settle prior to polymerization, which causes this material to behave as a bilayer. The ELM layer (100 μm thick), prepared with 10 HEA/0.1 BIS and 1 × 10^5^ CFU of ABEC per μL of ELM, was polymerized directly on top of the hydrogel layer, likely enabling monomer diffusion into the underlying layer and forming a strong interface between the two layers. CNCs swell, retain water, and enhance the mechanical properties of hydrogels when incorporated as reinforcing agents [74,75]. The hydrogel bilayer globally swells significantly more than the ELM, with volume increase of 192 ± 3% (for hydrogel layer) and 60 ± 2% (for ELM layer) (**Figure 7B**). Furthermore, swelling of the ELM leads to bending with the CNC-rich layer on the outside, indicating the CNC rich layer swells more. The apparent compressive modulus of the hydrogel bilayer (1.27 ± 0.16 MPa) is substantially greater than that of the ELM layer (13 ± 1 kPa) (**Figure 7C, Figure S14**). This difference allows the hydrogel to dominate bending. As the ELM is polymerized onto the CNC-poor side of the hydrogel, it always forms the inner face of the ELM-device after bending.

**Figure 7.**
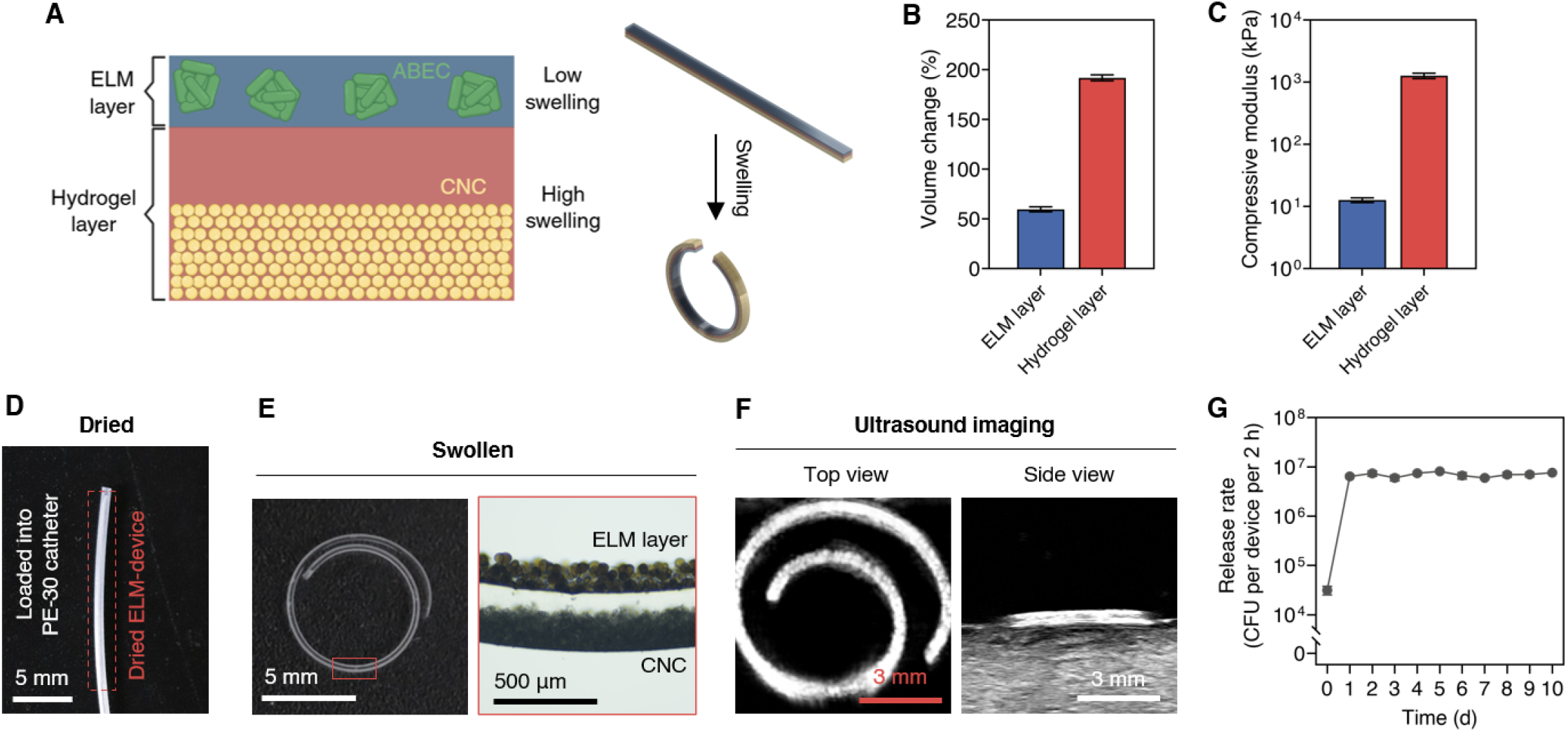
Shape-changing ELM-devices release ABEC in a sustained manner. **(A)** Schematic illustrating the bending mechanism of the ELM-device. **(B)** Volume increase of the different ELM-device layers (ELM and hydrogel layer) after swelling in PBS for 24 h. **(C)** Compressive modulus of the different ELM-device layers (ELM and hydrogel layer). **(D)** Photograph of a dried ELM-device loaded into a PE-30 catheter. **(E)** Photograph of a swollen ELM-device demonstrating bending, alongside an optical microscopy image showing a close-up view of the swollen ELM-device displaying the ELM and hydrogel layers (CNC-loaded and non-CNC-loaded regions). **(F)** Ultrasound images showing swollen and bent ELM-device. **(G)** ABEC release rate from ELM-device in human urine as a function of time. The ELM layer was prepared with ∼1 × 10^5^ CFU of ABEC per μL of ELM and incubated in 2 mL of human urine at 37 °C under aerobic conditions with shaking at 200 rpm. Media was refreshed every 24 h. Data in panels **(B)** and **(C)** are presented as mean ± standard deviations (*n* = 3), and data in panel **(F)** are presented as mean ± standard error of the mean (*n* = 3). Trend lines are only intended to guide the eye.

Incubating freshly prepared ELM-devices in LB media for 24 h allows the ABEC to proliferate within the ELM-devices, initiating release. This 1-day growth period prior to implantation reduces the lag typically observed before ABEC release from ELM begins. After the 1-day growth, the ELM-devices were dried for 24 h to fix a flat form (**Figure S15**). The device becomes stiff enough to enable loading into a catheter for intravesical delivery in mice (**Figure 7D**). When placed in PBS, the dried ELM-devices rapidly absorb water, swell, and bend, reaching a radius of curvature of 2.7 mm within 30 s and stabilizing at 3 mm within 5 min (**Figure 7E**). At stable curvature, the ELM layer (145 ± 7 μm thick) is clearly positioned on the inner face, the CNC-loaded region of the hydrogel layer (184 ± 2 μm thick) on the outside, and the non-CNC-loaded hydrogel region (91 ± 6 μm thick) in the middle (**Figure 7E**). The CNCs also bestow the material with echogenicity [76]. As a result, the ELM-devices can be visualized using ultrasound (**Figure 7F**), which can be exploited to visualize the ELM-devices in the bladder over time. Finally, we confirm that the dried, 1-day-grown ELM-devices release ABEC at a rate of 7.03 ± 0.75 × 10^6^ CFUs per ELM-device per 2 h and sustain this release for at least 10 days (**Figure 7G**).

### 2.8. ELM-devices release probiotic ABEC in mice bladder

Shape-changing ELMs can be delivered into the mouse bladder and retained for at least 2 days. To demonstrate sustained ABEC release in vivo, the ELM-devices must first be retained in the bladder for a defined period. For bladder delivery, ELM-devices prepared and processed as described above in 2.7 were used. These ELM-devices were loaded into PE-30 catheters (ID = 432 μm, OD = 762 μm), and a blunt 30G stylet was then added to the distal end with a slight tolerance between the needle and the ELM-device. The needle was separated from the distal end of the catheter by a parafilm stopper (**Figure 8A**). The catheter, preloaded with the ELM-device, needle, and stopper, was then transurethrally advanced into the bladder, and the ELM-devices were deployed using the stylet (**Figure 8A**). Each mouse received two ELM-devices. After delivery, each mouse was imaged using ultrasound every 24 h for 3 days to observe the presence of ELM-devices in the bladder (**Figure 8C**). Four mice were tested; all four retained ELM-devices for the first 2 days, but only one retained the ELM-devices for 3 days (**Figure 8D, Figure S16**). Mouse micturition frequency is 20 to 30 times per 24 h [77], suggesting that all four mice retained the ELM-devices for at least 40 voiding cycles.

**Figure 8.**
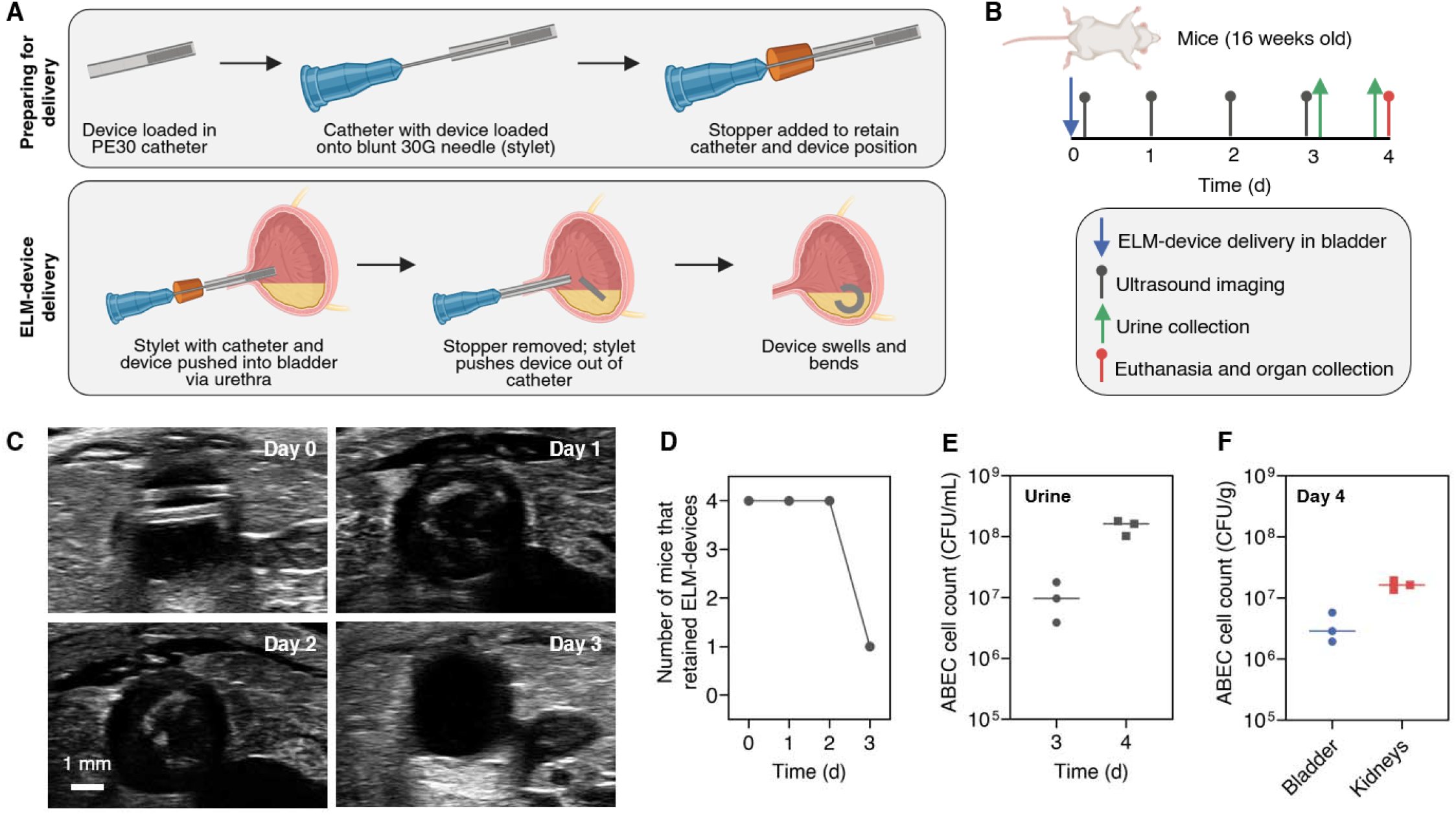
Shape-changing ELM-devices release ABEC in vivo. **(A)** Schematic illustrating the preparation of the ELM-device for delivery and the device delivery procedure in a mouse bladder. **(B)** Timeline showing the sequence of events for the in vivo study. **(C)** Ultrasound images of the mouse bladder captured for 3 days, showing ELM-device retention. **(D)** ELM-device retention in mice over 3 days. **(E)** ABEC counts in mouse urine 3 and 4 days after delivery. **(F)** ABEC counts in mouse bladder and mice kidneys 4 days after delivery. Four mice were implanted, with each mouse receiving two ELM-devices. All urine was collected prior to euthanasia, and all organs were collected following euthanasia. Data in **(E)** and **(F)** are presented as mean with individual data points (*n* = 3).

The ELM-devices release ABEC in the mouse bladder in a sustained manner. The mouse that retained the ELM-devices for 3 days was euthanized on day 3, and the ABEC counts in the bladder and kidneys were quantified as 5.8 × 10^7^ CFU/g and 9.8 × 10^7^ CFU/g, respectively (**Figure S17A**,**B**). It is important to note that sustained ABEC release in the bladder also leads to renal colonization, suggesting that ABEC could protect against both cystitis and pyelonephritis. ELM-devices were recovered from two mice and contained 2.14 ± 1.12 × 10^7^ CFU per ELM-device (**Figure S17C**), demonstrating that the ABEC loaded in the ELM-devices survived in the mouse bladder, utilizing nutrients from urine, for at least 3 days. Urine from the three mice that expelled their ELM-devices within 3 days were collected and quantified 3 and 4 days post-implantation. These mice have 1.05 ± 0.40 × 10^7^ CFU/mL of ABEC in urine on day 3 and 1.49 ± 0.23 × 10^8^ CFU/mL on day 4. All three mice were euthanized on day 4, and the ABEC counts in their bladder and kidneys were quantified. They have 3.54 ± 1.16 × 10^6^ CFU/g of ABEC in the bladder and 1.65 ± 0.17 × 10^7^ CFU/g of ABEC in the kidneys (**Figure 8F, Figure S18**). The persistence of ABEC in urine over multiple days could be attributed to sustained release from the ELM-devices, as transurethral inoculated free-floating ABEC do not persist in the mouse urinary tract [43]. The high ABEC counts (>10^6^ CFU/g) in both the bladder and kidneys further confirm sustained release from ELM-devices. Thus, using implantable ELMs, ABEC persistence in the mouse urinary tract can be achieved for at least 4 days. This approach may enable future studies testing the ability of ABEC to mitigate UPEC-induced UTI in a mouse model.

### 2.9. Limitations

In all in vitro experiments with urine refresh cycles, urine was refreshed manually using a pipette, which does not accurately reproduce physiological urine flow [78,79]. Under more realistic flow conditions, we anticipate lower retention of both free-floating and ELM-released ABEC [43], which may reduce the effectiveness of free-floating ABEC and may extend the time required for ELMs to achieve comparable reductions in UPEC load. In vitro competition and invasion assays were conducted over relatively short timescales (6 h), whereas clinical application would likely require demonstration of efficacy over longer periods. Because UPEC reduce urothelial cell viability [80–82], we were unable to extend these experiments further. In future work, we plan to evaluate ELM performance under continuous, physiologically relevant urine flow and over extended timescales using bladder organoids [83], which will provide a more accurate assessment of ELM efficacy against UTIs. Although we tested the effects of ABEC ELMs against two prototypical UPEC isolates (CFT073 and UTI89), future investigations could evaluate its effect against a broader range of UPEC isolates. The in vivo study was limited in scope with a small number of animals, and device retention over the full 3-days was achieved in only one mouse. A 3-day retention period may also be insufficient for prophylactic applications [84]. Moreover, here we only assessed ABEC persistence rather than prophylactic efficacy against UPEC infection. Finally, the presence of ABEC in the kidneys, while consistent with sustained bladder release, warrants further evaluation to confirm that ABEC colonization at these sites remains asymptomatic and safe over longer durations. The larger diameter of the ELM-device, relative to the capacity of a mouse bladder, may lead to contact with the bladder wall and induce inflammation [85]. The stiffness of the ELM-devices may decrease over time due to the fracture-based ABEC release mechanism [52]. Both factors could reduce the retention of shape-changing ELM-devices within the bladder. We aim to address these issues through refined device geometry, mechanical tuning, and optimized deployment to ensure consistent long-term residence in murine models. Building on the demonstrated in vivo ABEC persistence, subsequent work will test the ability of ABEC-releasing ELM-devices to control UPEC and prevent rUTI in murine models over multiple days. A comprehensive assessment of the safety, biocompatibility, and long-term host response to implanted ELMs, including confirmation that ABEC persistence in the bladder and kidneys remains asymptomatic, will also be critical. In addition, we plan to extend this approach to larger animal models, such as pigs that are naturally prone to developing rUTIs [86].

## 3. CONCLUSION

In this work, we developed and evaluated ELMs that release the urinary probiotic ABEC in a sustained manner as an innovative prevention strategy against rUTIs. Using in vitro models incorporating human urothelial cells, human urine, and periodic urine exchange to better mimic the bladder environment, we demonstrated that ABEC-releasing ELMs suppress UPEC proliferation and inhibit UPEC attachment to and invasion of urothelial cells. Because ELMs continuously release ABEC, they maintain competitive pressure against UPEC even as planktonic bacteria are cleared during voiding or refresh cycles, allowing them to outperform a single administration of free-floating ABEC and to remain effective at challenging ABEC-to-UPEC ratios. We further showed that UPEC colonization of the ELM is negligible, indicating these ELMs continue to release ABEC in a setting emulating UTI. In a rUTI model, sustained ABEC release from ELMs reduced the proliferation, re-adherence and re-invasion of UPEC expelled from previously infected urothelial cells. Finally, we designed shape-changing ELMs capable of transurethral delivery and retention within the murine bladder, and demonstrated sustained ABEC release in vivo, with ABEC persisting in the bladder, kidneys, and urine for at least 4 days. Collectively, these findings establish a framework for achieving sustained probiotic persistence at hard-to-access sites and highlight ELM-based implants as a promising antibiotic-free platform for the prevention of rUTIs.

## 4. METHODS

### 4.1. Materials

2-Hydroxyethyl acrylate (HEA), *N,N*′-methylenebis(acrylamide) (BIS), lithium phenyl-2,4,6-trimethylbenzoylphosphinate (LAP) were purchased from Sigma-Aldrich. Cellulose nanocrystals (CNCs) were purchased from CelluForce. Luria–Bertani (LB) Lennox broth and agar were purchased from BD Difco. Phosphate buffered saline (PBS) was purchased from Research Products International (RPI). PBS, LB broth, LB-agar were prepared with diH_2_O and were sterilized by autoclaving at 120 °C for 20 min and then stored at room temperature (PBS and LB broth) or at 4 °C (LB agar). Antibiotics were added at the following concentrations when required: kanamycin, 50 μg/mL; chloramphenicol, 32 μg/mL; and rifampin, 100 μg/mL. Dulbecco’s Phosphate-Buffered Saline (DPBS) and Roswell Park Memorial Institute (RPMI) 1640 media were purchased from Gibco (Thermo Fisher). Both DPBS and RPMI were stored at 4 °C and were warmed to 37 °C in a water bath before use. Filter-sterilized human urine that was pooled from 10 adult female donors who had no history of UTI or diabetes was purchased from Cone Bioproducts. Artificial urine was prepared by following the procedure given in [87], filter-sterilized and stored at 4 °C.

Bacterial strains ABEC Rif (Rifampin resistant ABEC), CFT073 Kan (Kanamycin resistant *E. coli* CFT073), UTI89 Kan (Kanamycin resistant *E. coli* UTI89) [39,43], and CFT073 Δ*hly*::Cam (CFT073 lacking hemolysin and chloramphenicol resistant) were used from the Subash Lab collection.

### 4.2. Bacterial cultures

Bacterial cultures were grown in 25 ml LB broth at 37 °C for 24 h at 200 rpm (*n* = 3), inoculated with single colonies from plain or antibiotic-containing LB agar plates, depending on the strain. Cultures of UTI89-Kan were grown under static conditions to induce production of type 1 fimbriae [88]. The cultures were centrifuged at 4000 rpm for 10 min at room temperature, bacterial cell pellets were then washed thrice with 25 mL of PBS, and the suspension was then diluted and adjusted to an optical density of 3.0 at 600 nm using a UV-vis spectrophotometer (Genesys 40, Thermo Scientific). Viable cell counts were obtained by plating serial dilutions on plain or antibiotic-containing LB agar plates using an automatic plater (easySpiral, Interscience). The plates were incubated at 37 °C overnight, and CFUs were counted using an automatic colony counter (Scan 300, Interscience). The bacterial suspensions had ∼2 × 10^9^ CFU/mL, which were then further diluted for preparing ELMs.

### 4.3. Preparation of ELMs

The ELMs were prepared by free radical polymerization of HEA (monomer) and BIS (crosslinker). LAP was used as a photoinitiator for this polymerization. Stock solutions of BIS (0.02 g/mL) and LAP (0.02 g/mL) were prepared in diH_2_O. All chemicals and stock solutions were filter sterilized (polyethersulfone, 0.2 μm) before ELM preparation. ELMs were prepared with 1 × 10^5^ cells/μL. The ELMs (10 HEA / 0.1 BIS) were prepared using 10 wt% HEA and 0.1 wt% BIS. The concentration of each component denotes the percentage by weight in the entire pre-gel solution, including cells. All ELMs were prepared with 0.04 wt% LAP. The remaining fraction of the pre-gel solution was filled with an equivalent mass of LB media. All the solutions were then mixed thoroughly using a vortex mixer.

After preparation, these pre-gel solutions were filled into polyethylene tubing (PE-50 (ID = 584 μm, OD = 965 μm), Braintree Scientific) and exposed to UV irradiation (UVP Crosslinker CL-3000, Analytik Jena) of 365 nm at an intensity of 1.2 mW/cm^2^ for 2 min to polymerize. The polymerized ELMs were then dispensed from the tubing with a sterile 27G needle, trimmed to a 4 mm length with razor blades, and washed thrice with PBS to remove unpolymerized monomer residues. The volume of the resulting ELMs was ∼1 μL.

### 4.4. Quantification of ABEC release from ELMs

To quantify the release of ABEC from ELMs, freshly prepared ELMs were placed in 14 mL round-bottom tubes containing 2 mL of media (LB medium or human urine or synthetic urine or PBS) and incubated at 37 °C under aerobic conditions with constant shaking at 200 rpm. After 2 h of incubation, an aliquot of the medium was serially diluted (10-fold) and plated on LB-Rif agar. Every 24 h, the ELMs were removed from the medium, washed three times with PBS, transferred to 2 mL of fresh media (LB medium or human urine or synthetic urine or PBS), and incubated under the same conditions. After 2 h of incubation, cell release was measured following the same procedure. The release of *E. coli* from the ELMs was measured for 10 days across all conditions.

### 4.5. Quantification of UPEC colonization of ELMs

Freshly prepared ELMs were first incubated in 2 mL of human urine for 3 days at 37 °C under aerobic conditions with constant shaking at 200 rpm, with ELMs washed and the urine refreshed every 24 h. After 3 days, the ELMs were washed three times with PBS, transferred to 2 mL of human urine supplemented with UPEC (CFT073-Kan) at 1 × 10^6^ cells/mL, and incubated at 37 °C under aerobic conditions with constant shaking at 200 rpm for 24 h. The ELMs were subsequently washed three times with PBS and placed in 3 mL of PBS for 30 min. Each ELM was then homogenized using a hand-held homogenizer (Homogenizer 150, Fisher) at high speed for approximately 30 s, until no visible hydrogel fragments remained. The homogenizer blade was decontaminated with 70% ethanol and rinsed with sterile diH_2_O between samples. Following homogenization, aliquots of the resulting suspension were serially diluted, plated on antibiotic-containing LB-agar plates, and colony-forming units (CFUs) were enumerated to determine the ABEC and UPEC counts.

UPEC infiltration into cell-free hydrogels and 3-day grown ELMs (dead) were tested following the same procedure. For cell-free hydrogels, freshly prepared cell-free hydrogels were washed three times with PBS and added to urine supplemented with CFT073-Kan. For 3-day grown ELMs (dead), 3-day grown living ELMs were exposed to 10 mL of 70% ethanol for 24 h, washed three times with PBS, and then added to urine supplemented with UPEC. All subsequent steps were the same as described above.

### 4.6. Competition experiments

Human urinary bladder epithelial cells, 5637 (ATCC, HTB-9) were grown in RPMI supplemented with 10% FBS and 1% PSG. The urothelial cells were seeded in 24-well plates at 4 × 10^4^ cells/well and incubated at 37 °C with 5% CO_2_ until reaching 80–90% confluency. The cells were then washed three times with 500 μL of DPBS, after which 500 μL of human urine was added to each well.

Three conditions were tested: (1) ELM, (2) free-floating, and (3) control. For the ELM condition, one 3-day-grown ELM was added per well. After 2 h, different concentrations (1 × 10^4^, 1 × 10^5^, 1 × 10^6^, 1 × 10^7^ CFU/mL) of UPEC (CFT073 Δ*hly*::Cam) were added to all conditions, corresponding to ABEC-to-UPEC ratios of 100:1, 10:1, 1:1, and 1:10. These ratios were based on the ∼1 × 10^6^ CFU/mL of ABEC released by ELMs within the first 2 h. For the free-floating condition, 1 × 10^6^ CFU/mL of ABEC was added simultaneously with UPEC to enable direct comparison with the ELM condition. The cells were then incubated for 6 h. Every 2 h, human urine was aspirated from the wells, and fresh human urine was added. The removed supernatant was diluted and plated on LB-Rif and LB-Cam agar plates to quantify ABEC and UPEC counts, respectively.

### 4.7. Quantification of bacterial attachment to and invasion into urothelial cells

After 6 h of competition, ABEC and UPEC that had attached to or invaded the urothelial cells were quantified by washing the urothelial cells three times with DPBS, and lysing with 0.1% Triton X-100. The cells were then manually scraped using a 1000 μL sterile pipette tip, collected, serially diluted, and plated on LB-Rif and LB-Cam agar plates to quantify the ABEC and UPEC counts, respectively. The bacterial counts obtained from the lysate represent both attached and intracellular bacteria.

To measure intracellular ABEC and UPEC counts specifically, after 6 h of competition, the cells were washed three times with DPBS and treated with 1 mL of gentamicin (100 mg/mL) for 2 h to kill all extracellular bacteria. The gentamicin was then removed, and the cells were washed three times with 1 mL of DPBS. The urothelial cells were subsequently lysed with 0.1% Triton X-100, and processed as described above to quantify intracellular ABEC and UPEC counts. Adherent bacterial counts were determined by subtracting the intracellular counts from the total counts obtained from the lysate.

### 4.8. Evaluation of ABEC against UPEC-infected urothelial cells

Urothelial cells were seeded in 24-well plates and grown as described in Section 4.6. The cells were then infected with UPEC (UTI89-Kan) at an MOI of 100. The plates were centrifuged at 500 rpm for 5 min and incubated for 30 min. The cells were washed three times with DPBS to remove non-adherent bacteria and then treated with 100 mg/mL gentamicin for 2 h to kill all extracellular bacteria. The cells were subsequently washed and lysed as described in 4.7 to determine intracellular bacterial counts at 0 h (T_o_).

Following gentamicin treatment and washing, 500 μL of human urine was added to each well. Three conditions were tested: (1) negative control, (2) positive control, and (3) free-floating. For the negative control, only urine was added. For positive control, the urine was supplemented with 100 mM methyl α-D-mannoside and 25 μg/mL trimethoprim (a bacteriostatic antibiotic). For the free-floating condition, 1 × 10^6^ CFU/mL of ABEC was added to the urine. All co-cultures were incubated for 6 h under static conditions, with no urine replacement during the experiment. After 6 h, the urine (supernatant) was collected from each well, diluted, and plated on appropriate antibiotic-containing LB agar plates to quantify ABEC and UPEC counts. Total cell-associated ABEC and UPEC (attached to and intracellular) were also quantified after 6 h, essentially as described above in 4.7. To quantify bacterial counts in the supernatant and lysate at 2 h, 4 h, and 6 h, the same procedure was followed for the negative control and free-floating conditions, with supernatants and lysates collected and plated at each timepoint.

### 4.9. Evaluation of ELM against UPEC-infected urothelial cells

Urothelial cells in 24-well plates were infected with UPEC (UTI89-Kan), lysed and intracellular bacterial counts at 0 h (T_o_) were determined following the same procedure described in Section 4.8. After 500 μL of human urine was added to each well, two conditions were tested: (1) control and (2) ELM. For the control, only urine was added. For the ELM condition, one 3-day-grown ELM, which releases ∼1 × 10^6^ CFU/mL of ABEC within 2 h, was added per well. Both conditions were incubated under static conditions for 6 h. Every 2 h, human urine was manually aspirated from the wells and replaced with fresh urine. The removed supernatant was diluted and plated to determine ABEC and UPEC CFUs. After 6 h, ABEC and UPEC that had attached to or invaded the urothelial cells were quantified as described in 4.7.

### 4.10. Preparation of precursors for the ELM-devices

The ELM-device consists of two layers prepared from two distinct precursor solutions. The ELM layer precursor was prepared by mixing 10 wt% HEA, 0.1 wt% BIS, and 0.04 wt% LAP with 1 × 10_ CFU/μL of ABEC, where each component concentration represents its weight percentage in the entire pre-gel solution, including cells. The remaining fraction of the solution was filled with an equivalent mass of LB medium. BIS and LAP were added from stock solutions (0.02 g/mL each) prepared in diH_2_O. All components were thoroughly mixed using a vortex mixer to obtain the final ELM layer precursor. HEA, BIS, and LAP stock solutions were filter-sterilized (polyethersulfone, 0.2 μm) prior to ELM preparation.

The hydrogel layer precursor was prepared by directly combining 2 wt% BIS, 69 wt% HEA, and 2 wt% AM, followed by vortexing and brief heating (3–6 s) with constant swirling to ensure dissolution. The solution was then cooled to room temperature, after which 25 wt% CNC and 0.04 wt% LAP were added and vortexed. LAP was added from a stock solution (0.02 g/mL) prepared in diH_2_O. The resulting mixture served as the hydrogel layer precursor. Both precursor solutions were subsequently used for ELM-device preparation.

### 4.11. Preparation of the ELM-devices

Three glass slides (75 mm × 50 mm × 1 mm) were used, two of which were treated with a water-repellent spray (Rain-X Original). A treated and an untreated glass slide were separated by a 175 μm spacer and secured with clips, with the treated slide positioned on top and the untreated slide on the bottom. The hydrogel layer precursor was then filled into the mold and left to rest for 1 min, allowing the CNCs to settle onto the bottom slide and form a distinct layer within the hydrogel. The mold was subsequently exposed to UV irradiation (same conditions described in Section 4.3) for 1 min.

The top (treated) glass slide was then removed, and an additional 100 μm spacer was added on top of the existing 175 μm spacer, bringing the total thickness to 275 μm. A new treated glass slide was placed on top and secured with clips. The ELM layer precursor was filled into the mold and exposed to UV irradiation for 30 s, after which the mold was flipped and irradiated for another 1 min, and flipped again and irradiated for another 30 s. The top treated glass slide was then removed, unpolymerized monomers were washed away with PBS, and the bilayer was cut using two blades placed in parallel, where the gap between the blade edges defined the ELM-device width and the blade length (25 mm) defined the ELM-device length.

Each ELM-device was washed with PBS and then placed in a 14 mL round-bottom tube containing 2 mL of LB medium and incubated at 37 °C under aerobic conditions with constant shaking at 200 rpm. After 24 h, the ELM-devices were washed with PBS and dried on polytetrafluoroethylene (PTFE) sheets for 24 h at room temperature (∼20 °C) with a relative humidity of ∼30–50%. The dried ELM-devices were loaded into a polyethylene catheter (PE30, Braintree Scientific) using tweezers and a blunt 30G needle as a stylet, positioning each ELM-device near the proximal end while inserting the needle into the distal end with a slight tolerance between the needle and the ELM-device. A parafilm stopper was applied at the needle-catheter junction at the distal end to maintain a gap between the needle tip and the ELM-device.

### 4.12. Swelling studies

Freshly prepared 1 mm thick hydrogels corresponding to the hydrogel layer (69 HEA/2.0 AM/2.0 BIS/25 CNC) and ELM layer (10 HEA/0.1 BIS, without ABEC) of the ELM-device were washed with PBS and punched into 6 mm diameter discs. Immediately after washing, the front and top views of the hydrogels were photographed using a DSLR camera, and the images were analyzed using ImageJ to determine the initial volume. These hydrogels were then immersed in PBS for 24 h to reach swelling equilibrium, after which they were imaged and analyzed again to determine the swollen volume and calculate the change in volume for each sample.

### 4.13. Mechanical characterization

Cell-free hydrogel samples (2 mm thick) prepared with 10 HEA/0.1 BIS (corresponding to the ELM layer of the device) and 69 HEA/2.0 AM/2.0 BIS/25 CNC (corresponding to the hydrogel layer of the device) were equilibrated in PBS at room temperature for 24 h and cut into 12 mm diameter discs for characterization. Uniaxial compression tests were performed at room temperature using a dynamic mechanical analyzer (RSA-G2, TA Instruments). Each sample was loaded between the plates and immersed in ∼30 mL of PBS, then compressed at a strain rate of 0.05 mm/s. The compression modulus was calculated from the linear region of the stress-strain response, between 0.2% and 5% strain.

### 4.14. ELM-device imaging and bending measurements

Dried 1-day-grown ELM-devices were placed in 10 mL of PBS, and top-view images were captured using a digital single-lens reflex (DSLR) camera (EOS Rebel T7i, Canon) at 30 s, 1 min, 5 min, and 24 h. The images were processed in ImageJ to quantify the radius of curvature.

For thickness measurements, dried 1-day-grown ELM-devices were reswollen in PBS for 24 h and imaged from the top using a polarized optical microscope (Eclipse LV100N POL, Nikon) in transmission mode. The thickness of each layer (the hydrogel layer, including the CNC- and non-CNC-loaded regions, and the ELM layer) was measured using the distance measurement tool in the NIS-Elements software.

### 4.15. In vitro ultrasound imaging

Dried ELM-devices were immersed in PBS and allowed to swell for 5 min prior to imaging. Ultrasound imaging was performed using the Vevo F2 imaging system (Fujifilm VisualSonics) with a UHF57X linear transducer operating at a center frequency of 38 MHz. The transducer was positioned in contact with the PBS, slightly above the ELM-device. 2D ultrasound images were acquired, and a 3D scan was performed using the system scanning feature to obtain top and side views of the ELM-devices.

### 4.16. Quantification of ABEC release from ELM-devices prepared for delivery

To quantify the release of ABEC from ELM-devices, dried 1-day-grown ELM-devices were placed in 14 mL round-bottom tubes containing 2 mL of human urine and incubated at 37 °C under aerobic conditions with constant shaking at 200 rpm. After 2 h of incubation, an aliquot of the medium was serially diluted and plated on LB-Rif agar plates. The plates were incubated at 37 °C overnight, and CFUs were counted. Every 24 h, the ELM-devices were removed from the medium, washed with PBS, transferred to 2 mL of fresh urine, and incubated under the same conditions. After 2 h of incubation, cell release was measured following the same procedure. *E. coli* release from the ELM-devices was measured for 10 days.

### 4.17. Mouse experiments

In vivo experiments described in this study were approved by the Institutional Animal Care and Use Committee at Texas A&M University (protocol #2024-0201). Five female CBA/J mice (Jackson Laboratories) of 16 weeks of age were used in experiments.

### 4.18. In vivo ultrasound imaging

Mice were anesthetized with isoflurane and placed on a heating pad (37 °C), with continuous isoflurane delivery through a nose cone to maintain anesthesia. Hair removal cream (Nair Body Cream) was applied to the lower abdomen, and hair was removed. The area was then cleansed with water, and ultrasound gel was applied. Ultrasound imaging was performed using the Vevo F2 imaging system with UHF57X and UHF71X linear transducers operating at center frequencies of 38 MHz and 48 MHz, respectively. 2D ultrasound images were acquired, and a 3D scan was performed using the system’s scanning feature to obtain top and side views of the ELM-devices. Each imaging session lasted ∼20 min per mouse, during which the animal remained on the heating pad under isoflurane anesthesia. Imaging was performed daily for 4 days.

### 4.19. ELM-device delivery

Mice were anesthetized with isoflurane, lubricant (OB Lube, Priority Care) was applied to the outer genitalia, and the catheter (preloaded with the ELM-device, stylet, and stopper) was inserted through the urethra into the bladder. The stopper was then removed with tweezers, and the stylet was fully advanced into the catheter five times to deploy the ELM-device into the bladder lumen. The catheter and stylet were then withdrawn from the urethra, and the procedure was repeated for the second ELM-device. Each mouse received two ELM-devices.

A total of five mice underwent implantation; however, only four received two ELM-devices each. The remaining mouse received a single ELM-device due to difficulties encountered during catheterization. This mouse was excluded from subsequent analysis.

### 4.20. Quantification of ABEC in vivo

Spontaneously voided urine was collected from each mouse individually, serially diluted and plated on LB-Rif agar. Following euthanasia, the bladder and kidneys were aseptically excised, placed in 3 mL of PBS, homogenized, serially diluted, and plated on LB-Rif agar. Plates were incubated overnight at 37 °C, and CFUs were enumerated. Viable counts were normalized to urine volume or tissue mass. ELM-devices recovered from the bladders were also homogenized and viable counts were determined as described above for organs.

### 4.21. Statistical analysis

Statistical analysis was performed using GraphPad Prism (Version 11.0.1). Data are shown as the mean ± standard error of means. For all statistical tests, *P* < 0.05 was set for statistical significance. Single comparisons were performed using a Mann-Whitney U test (unpaired), and multiple comparisons were performed using a Kruskal-Wallis test with a post-hoc Dunn’s test.

## ACKNOWLEDGEMENTS

Research reported in this publication was partially supported by the National Institute of Biomedical Imaging and Bioengineering of the National Institutes of Health under Award No. R56EB032395 (T.H.W., S.S., and P.E.Z.). This material is also partially based upon work supported by the National Institutes of Health under Grant No. DK142927 (S.S.). P.E.Z. receives endowment support from the Cain Foundation through the Felecia and John Cain Distinguished Chair in Women’s Health established in honor of Philippe E. Zimmern, MD. M.S.K. was supported through Texas A&M University Dissertation Fellowship. The Funders had no role in study design, data collection and analysis, decision to publish, or preparation of the manuscript. The content is solely the responsibility of the authors and does not necessarily represent the official views of the funders. Some of the graphics were created with BioRender.com.

