## Supporting Information for "Engineered living materials suppress uropathogenic *E. coli* growth and invasion of urothelial cells through sustained probiotic release"

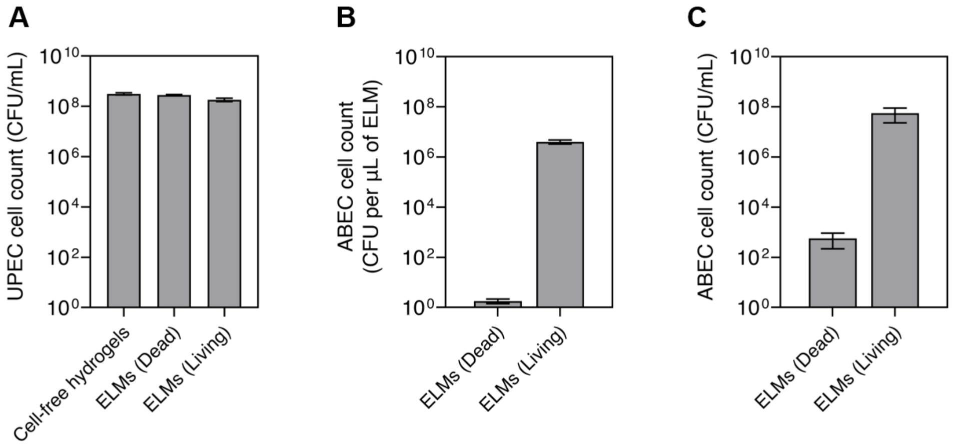


**Figure S1. Infiltration of UPECs into ELMs. (A)** UPEC counts in human urine containing ELMs or cell-free hydrogels after 24 h of incubation. **(B)** ABEC counts in human urine containing ELMs after 24 h of incubation. **(C)** ABEC counts within ELMs after 24 h of incubation with UPECs. Cell-free hydrogels and 3-day-grown ELMs (living and dead) were incubated in 2 mL of human urine supplemented with ~1 × $10^6$ CFU/mL of UPEC (CFT073-Kan) at 37 °C with shaking at 200 rpm under aerobic conditions for 24 h. Data are presented as mean ± standard error of the mean (*n* = 5).

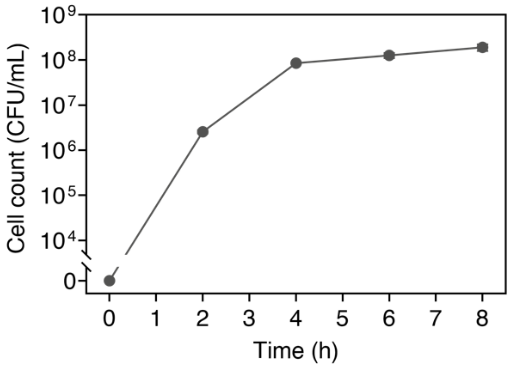


**Figure S2.** ABEC release from ELMs as a function of time in human urine in the presence of 5637 human urothelial cells. ELMs were incubated in 24-well plate at 37 °C with 5% $CO_2$, and human urine was refreshed every 2 h. Data are presented as mean ± standard error of the mean (*n* = 6). Trend lines are only intended to guide the eye.

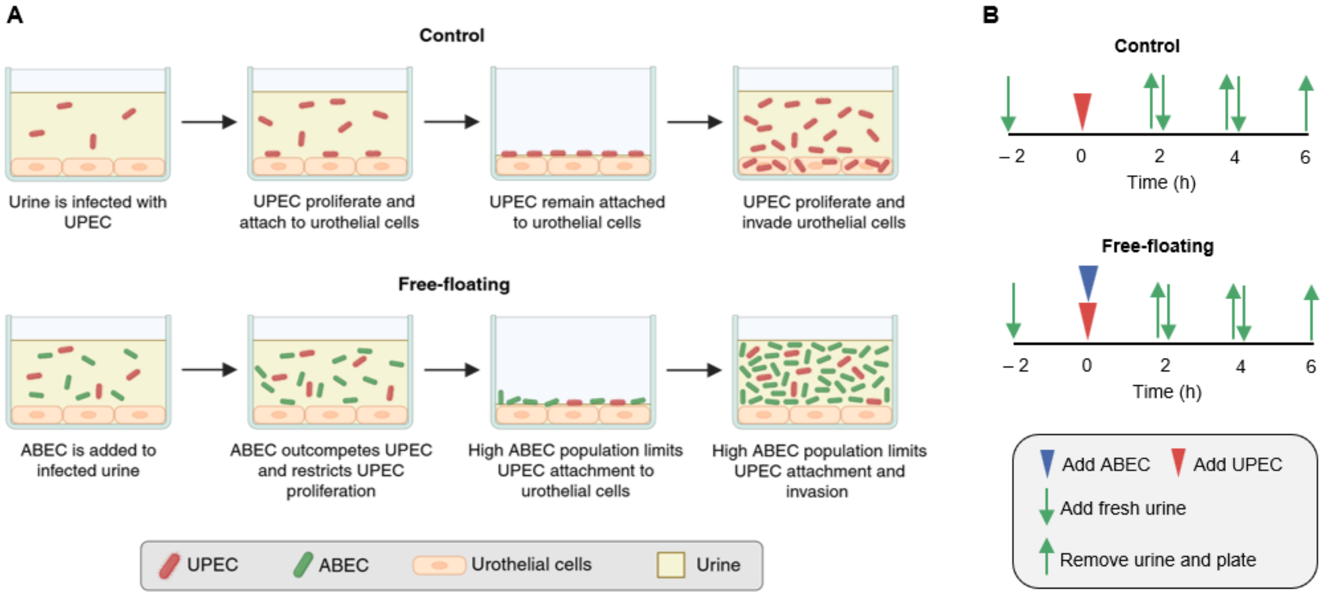


**Figure S3. ABEC restricts UPEC proliferation. (A)** Schematic illustrating UPEC proliferation in urine in the presence of urothelial cells, comparing conditions with and without free-floating ABEC. **(B)** Timeline of the competition experiment performed in 24-well plates in the presence of urothelial cells for control and free-floating conditions.

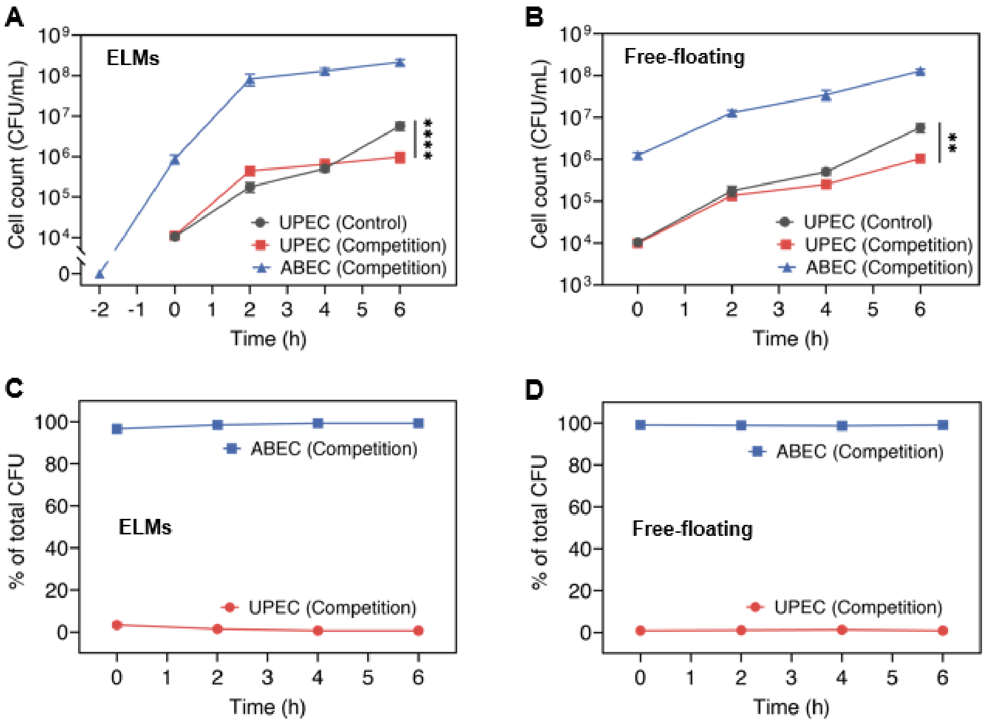


**Figure S4. ELMs and free-floating ABEC suppress UPEC proliferation at an initial ABEC:UPEC ratio of 100:1. (A, B)** ABEC and UPEC counts in the supernatant (urine) over 6 h of competition, comparing the control condition with the **(A)** ELM and **(B)** free-floating conditions. **(C, D)** Relative abundance of ABEC and UPEC in the supernatant in the **(C)** ELM and **(D)** free-floating conditions. Experiments were performed in 24-well plates incubated at 37 °C with 5% $CO_2$. ELMs were added at –2 h, UPEC and free-floating ABEC were introduced at 0 h, and human urine was refreshed every 2 h for 6 h. Data are presented as mean ± standard error of the mean ($n$ = 9). Statistical analysis in panel **(A)** and **(B)** was performed using a two-tailed Mann–Whitney $U$ test. * $P \leq 0.05$, ** $P \leq 0.01$, *** $P \leq 0.001$, **** $P \leq 0.0001$, and not significant (ns) for $P > 0.05$. Trend lines are only intended to guide the eye.

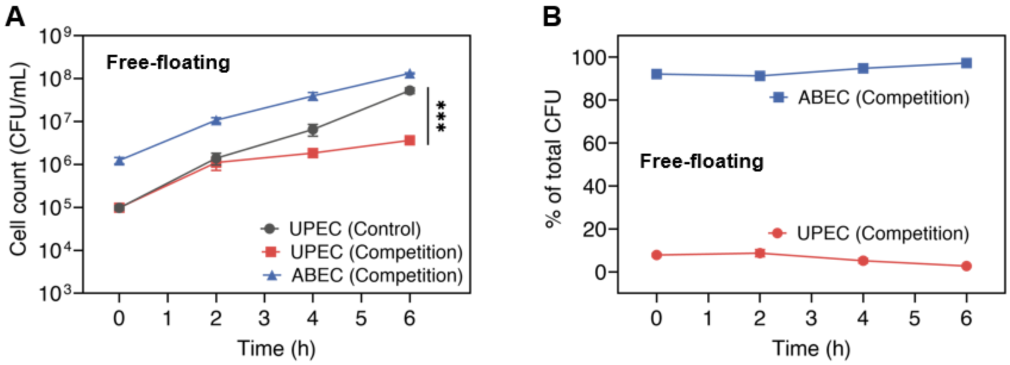


**Figure S5. Free-floating ABEC suppress UPEC proliferation at an initial ABEC:UPEC ratio of 10:1. (A)** ABEC and UPEC counts in the supernatant (urine) over 6 h of competition, comparing the control with the free-floating conditions. **(B)** Relative abundance of ABEC and UPEC in the supernatant in the free-floating condition. Experiments were performed in 24-well plates incubated at 37 °C with 5% $CO_2$. ELMs were added at –2 h, UPECs and free-floating ABEC were introduced at 0 h, and human urine was refreshed every 2 h for 6 h. Data are presented as mean ± standard error of the mean ($n$ = 9). Statistical analysis in panel **(A)** was performed using a two-tailed Mann–Whitney $U$ test. * $P \leq 0.05$, ** $P \leq 0.01$, *** $P \leq 0.001$, **** $P \leq 0.0001$, and not significant (ns) for $P > 0.05$. Trend lines are only intended to guide the eye.

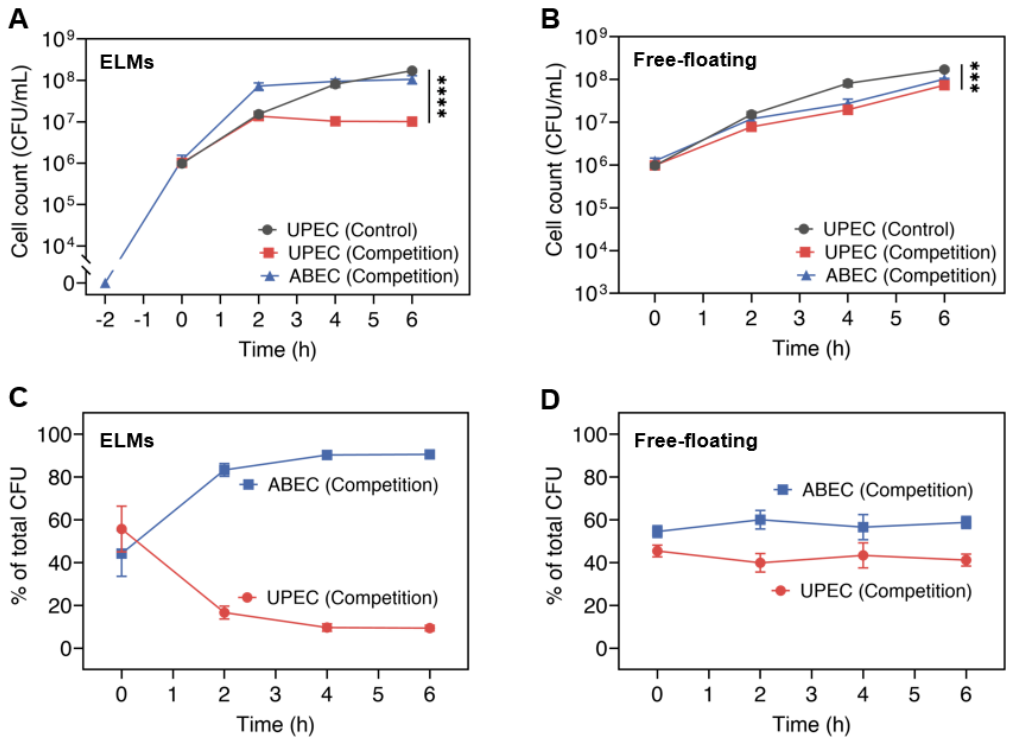


**Figure S6. ELMs and free-floating ABEC suppress UPEC proliferation at an initial ABEC:UPEC ratio of 1:1. (A, B)** ABEC and UPEC counts in the supernatant (urine) over 6 h of competition, comparing the control condition with the **(A)** ELM and **(B)** free-floating conditions. **(C, D)** Relative abundance of ABEC and UPEC in the supernatant in the **(C)** ELM and **(D)** free-floating conditions. Experiments were performed in 24-well plates incubated at 37 °C with 5% $CO_2$. ELMs were added at –2 h, UPEC and free-floating ABEC were introduced at 0 h, and human urine was refreshed every 2 h for 6 h. Data are presented as mean ± standard error of the mean ($n$ = 9). Statistical analysis in panel **(A)** and **(B)** was performed using a two-tailed Mann–Whitney $U$ test. * $P \leq 0.05$, ** $P \leq 0.01$, *** $P \leq 0.001$, **** $P \leq 0.0001$, and not significant (ns) for $P > 0.05$. Trend lines are only intended to guide the eye.

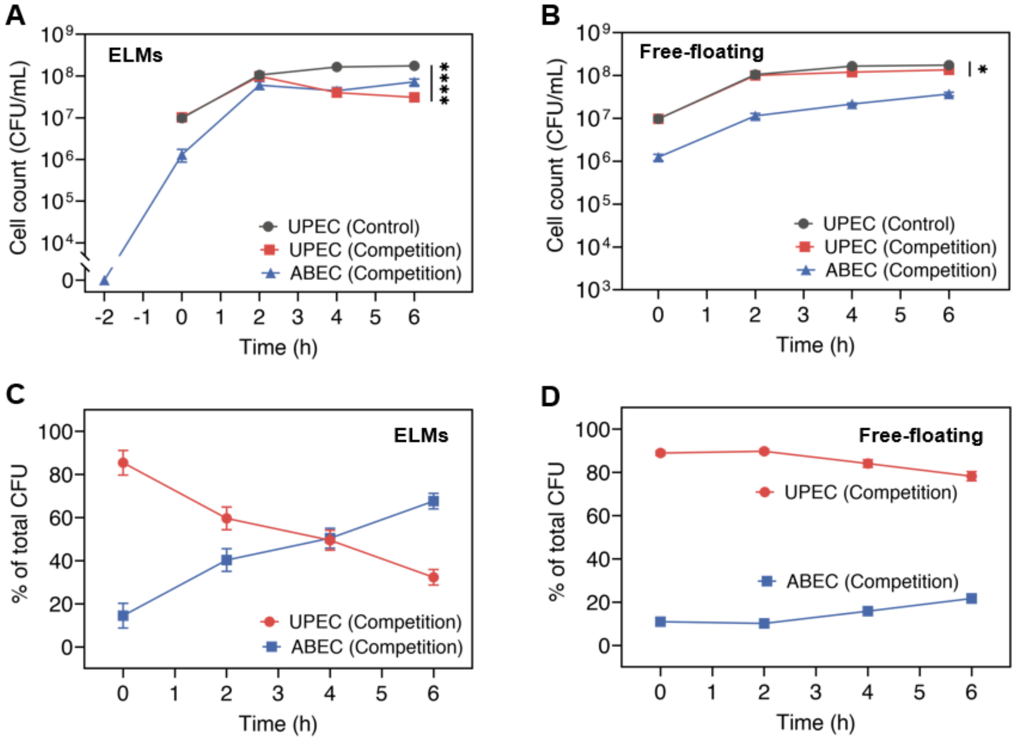


**Figure S7. ELMs and free-floating ABEC suppress UPEC proliferation at an initial ABEC:UPEC ratio of 1:10. (A, B)** ABEC and UPEC counts in the supernatant (urine) over 6 h of competition, comparing the control condition with the **(A)** ELM and **(B)** free-floating conditions. **(C, D)** Relative abundance of ABEC and UPEC in the supernatant in the **(C)** ELM and **(D)** free-floating conditions. Experiments were performed in 24-well plates incubated at 37 °C with 5% $CO_2$. ELMs were added at –2 h, UPEC and free-floating ABEC were introduced at 0 h, and human urine was refreshed every 2 h for 6 h. Data are presented as mean ± standard error of the mean ($n$ = 9). Statistical analysis in panel **(A)** and **(B)** was performed using a two-tailed Mann–Whitney $U$ test. * $P \leq 0.05$, ** $P \leq 0.01$, *** $P \leq 0.001$, **** $P \leq 0.0001$, and not significant (ns) for $P > 0.05$. Trend lines are only intended to guide the eye.

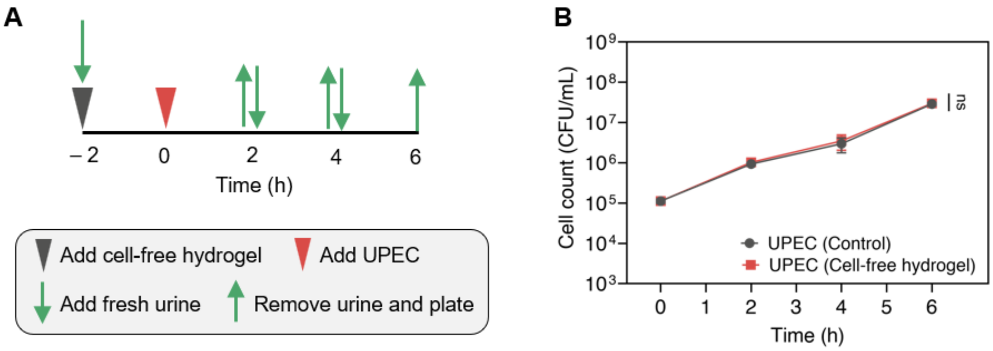


**Figure S8. Cell-free hydrogels do not reduce UPEC proliferation. (A)** Timeline of the experiment performed in the presence of urothelial cells for cell-free hydrogel condition. **(B)** UPEC counts in the supernatant (urine) over 6 h of competition in control and cell-free hydrogel conditions. Experiments were performed in 24-well plates incubated at 37 °C with 5% $CO_2$. Cell-free hydrogels were added at –2 h, UPEC were introduced at 0 h, and human urine was refreshed every 2 h for 6 h. Data are presented as mean ± standard error of the mean ($n$ = 3). Statistical analysis in panel **(B)** was performed using a two-tailed Mann–Whitney $U$ test. Not significant (ns) for $P > 0.05$. Trend lines are only intended to guide the eye.

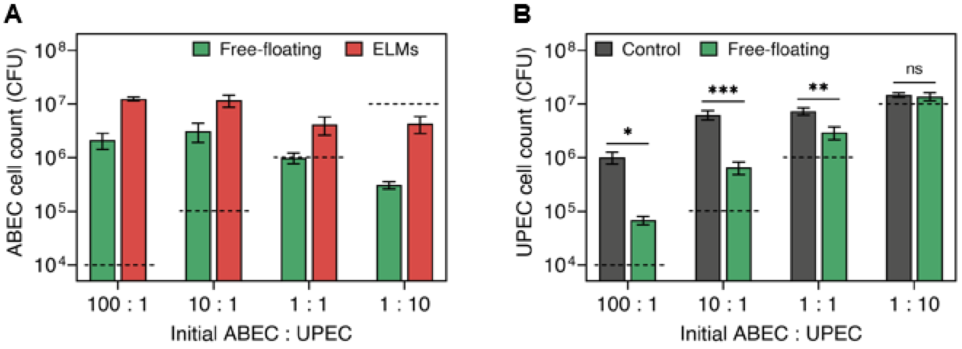


**Figure S9. High ABEC populations inhibit UPEC attachment to urothelial cells. (A)** ABEC counts in the lysate after 6 h of competition under free-floating and ELM conditions at different initial ABEC:UPEC ratios (100:1, 10:1, 1:1, and 1:10). **(B)** UPEC counts in the lysate (including both attached and intracellular UPECs) after 6 h of competition under free-floating conditions, compared to control conditions, at different initial ABEC:UPEC ratios (100:1, 10:1, 1:1, and 1:10). Dashed lines in panel **(A)** and **(B)** represent the initial UPEC concentration added at each ratio across all conditions (control, free-floating, and ELM). Experiments were performed in 24-well plates incubated at 37 °C with 5% $CO_2$. ELMs were added at –2 h, UPEC and free-floating ABEC were introduced at 0 h, and human urine was refreshed every 2 h for 6 h. Data are presented as mean ± standard error of the mean ($n$ = 9). Statistical analysis in panel **(B)** was performed using a two-tailed Mann–Whitney $U$ test. * $P \leq 0.05$, ** $P \leq 0.01$, *** $P \leq 0.001$, and not significant (ns) for $P > 0.05$.

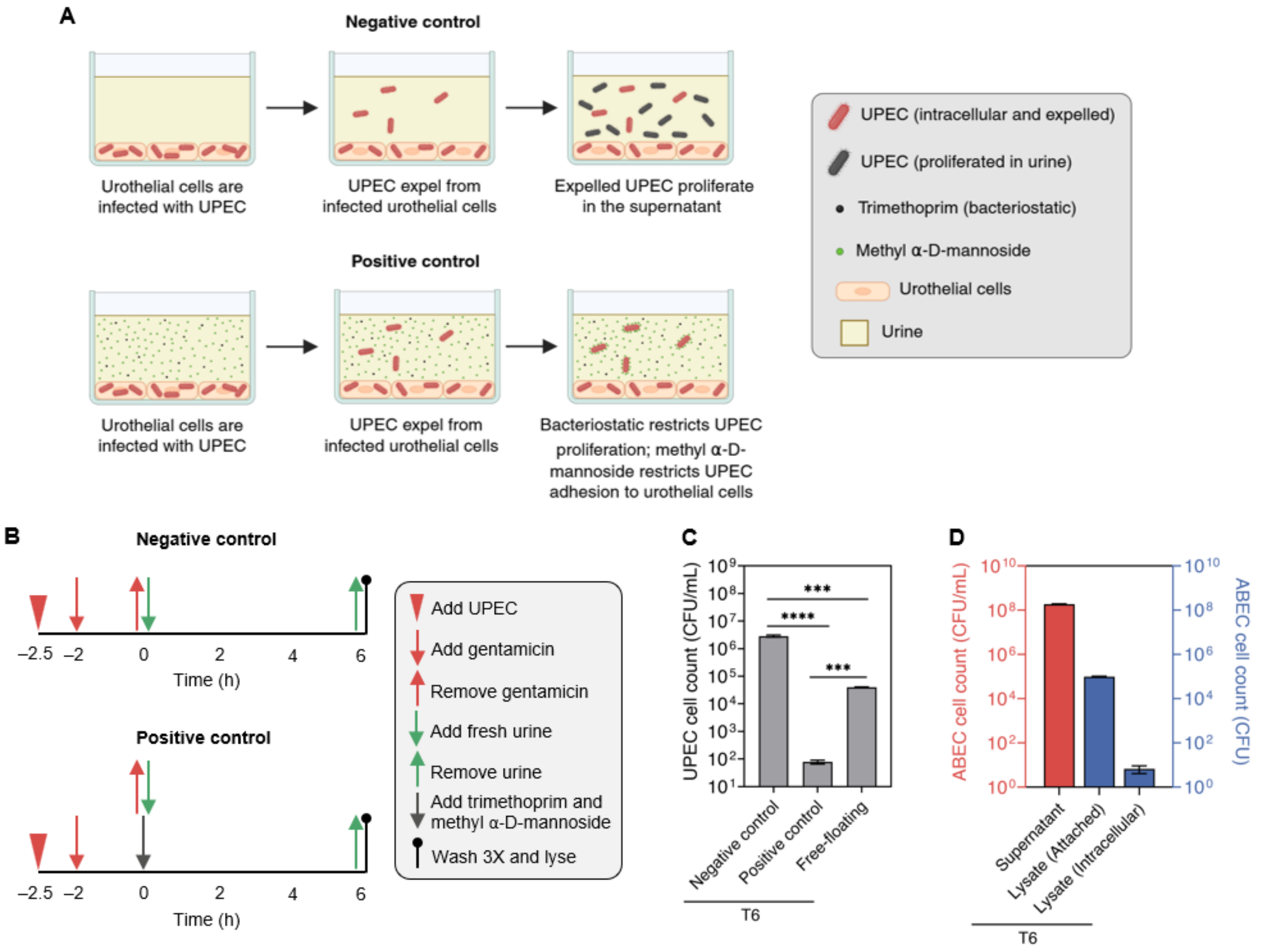


**Figure S10. ABEC inhibits proliferation of expelled UPECs. (A)** Schematic illustrating the expulsion of UPECs and their subsequent proliferation under negative control conditions, and their inability to proliferate under positive control conditions due to the presence of a bacteriostatic agent. **(B)** Timeline of the invasion experiment performed in the presence of urothelial cells for both (negative and positive) control conditions. **(C)** UPEC counts in the supernatant (urine) after 6 h of the invasion experiment under negative control, positive control, and free-floating conditions. **(D)** ABEC counts in the supernatant and lysate after 6 h of the invasion experiment under free-floating conditions. Experiments were performed in 24-well plates incubated at 37 °C with 5% $CO_2$. Free-floating ABEC (free-floating condition) or trimethoprim with methyl α-D-mannoside (positive control condition) were added post-invasion, and human urine was not refreshed for 6 h in either case. Data are presented as mean ± standard error of the mean ($n$ = 10). Statistical analysis in panel **(C)** was performed using a Kruskal–Wallis test with Dunn's multiple comparisons test. *** $P \leq 0.001$, **** $P \leq 0.0001$, and not significant (ns) for $P > 0.05$.

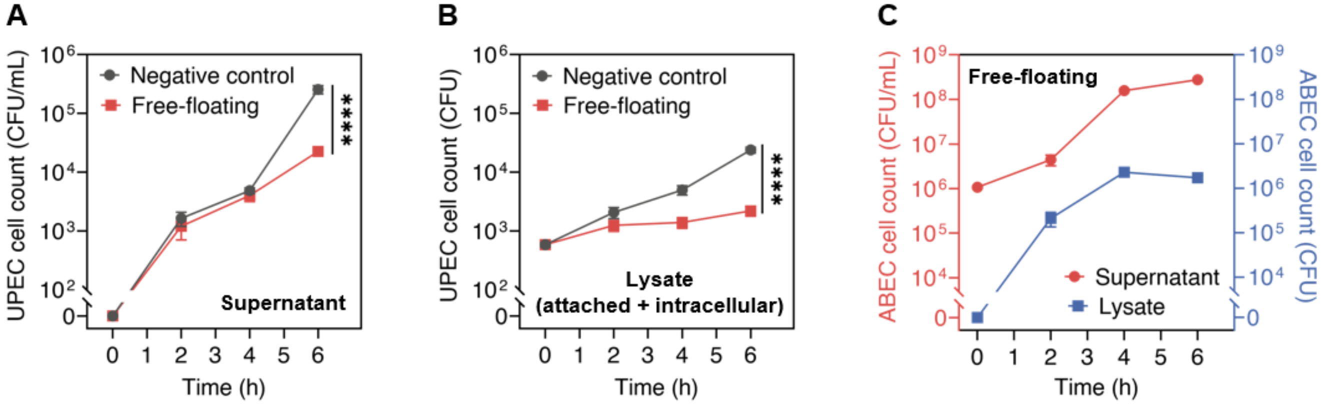


**Figure S11. ABEC inhibits proliferation of expelled UPECs. (A)** UPEC counts in the supernatant (urine) over the 6 h invasion experiment under free-floating conditions, compared to the negative control. **(B)** UPEC counts in the lysate (including both attached and intracellular UPEC) over the 6 h invasion experiment under free-floating conditions, compared to the negative control. **(C)** ABEC counts in the supernatant and lysate over the 6 h invasion experiment under free-floating conditions. Experiments were performed in 24-well plates incubated at 37 °C with 5% $CO_2$. Free-floating ABEC were added post-invasion, and human urine was not refreshed for 6 h. Data are presented as mean ± standard error of the mean ($n$ = 9). Statistical analysis in panel **(B)** and **(C)** was performed using a two-tailed Mann–Whitney $U$ test. **** $P \leq 0.0001$ and not significant (ns) for $P > 0.05$. Trend lines are only intended to guide the eye.

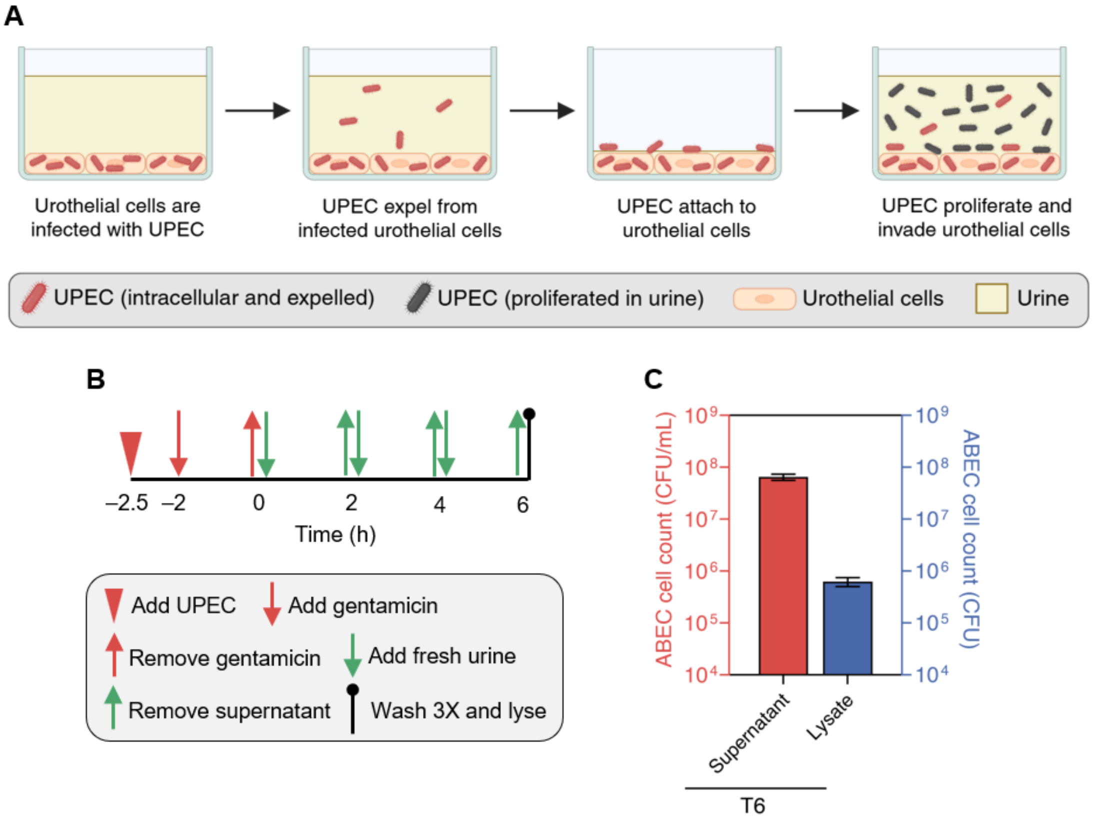


**Figure S12. ELMs inhibit proliferation of expelled UPECs. (A)** Schematic illustrating the expulsion of UPEC and the subsequent proliferation of the expelled UPEC under control conditions. **(B)** Timeline of the invasion experiment performed in the presence of urothelial cells under control conditions. **(C)** ABEC counts in the supernatant and lysate after 6 h of invasion experiment in the ELM condition. Experiments were performed in 24-well plates incubated at 37 °C with 5% $CO_2$. ELMs were added post-invasion, and human urine was refreshed every 2 h for 6 h. Data are presented as mean ± standard error of the mean ($n$ = 10).

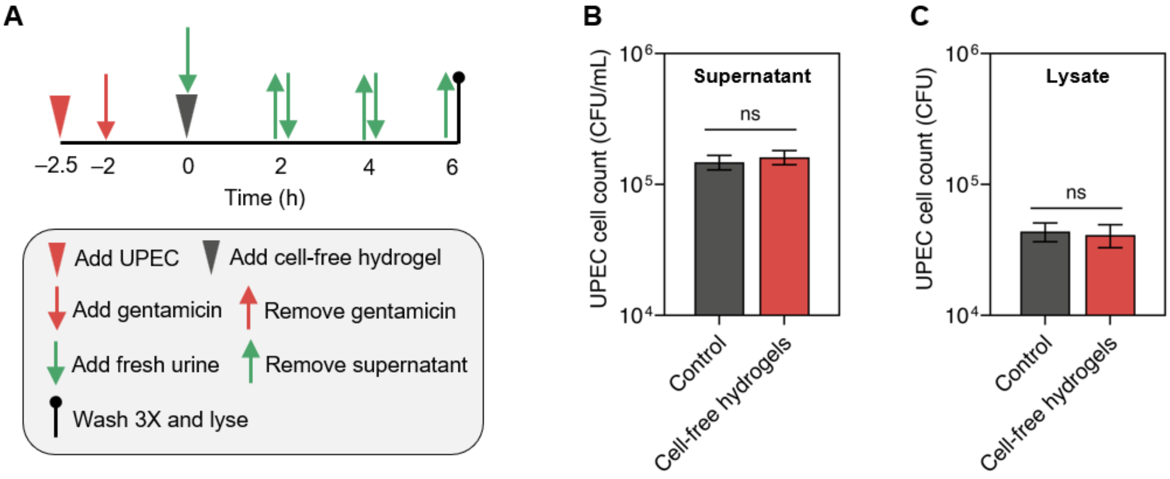


**Figure S13. Cell-free hydrogels do not reduce proliferation of expelled UPEC. (A)** Timeline of the invasion experiment performed in the presence of urothelial cells for the cell-free hydrogel condition. **(B)** UPEC counts in the supernatant (urine) after 6 h of the invasion experiment in the cell-free hydrogel condition compared to the control condition. **(C)** UPEC counts in the lysate (including both attached and intracellular UPEC) after 6 h of the invasion experiment in the cell-free hydrogel condition compared to the control condition. Experiments were performed in 24-well plates incubated at 37 °C with 5% $CO_2$. Cell-free hydrogels were added post-invasion, and human urine was refreshed every 2 h for 6 h. Data are presented as mean ± standard error of the mean ($n$ = 10). Statistical analysis in panel **(B)** and **(C)** was performed using a two-tailed Mann–Whitney $U$ test. Not significant (ns) for $P > 0.05$.

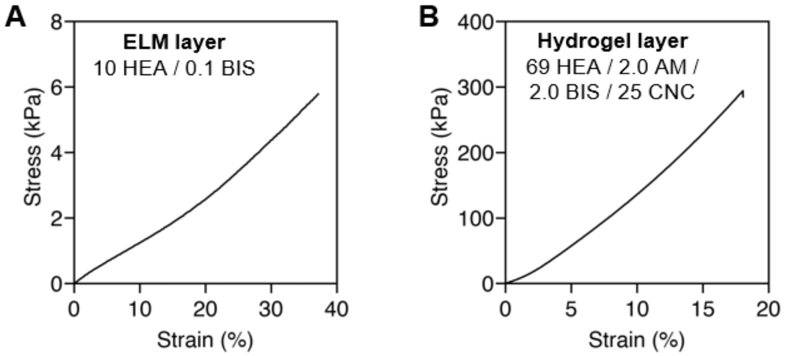


**Figure S14.** Stress-strain curves of different ELM-device layers (*n* = 3). **(A)** ELM layer (10 HEA / 0.1 BIS, no bacteria). **(B)** Hydrogel layer (69 HEA / 2 AM / 2 BIS / 25 CNC).

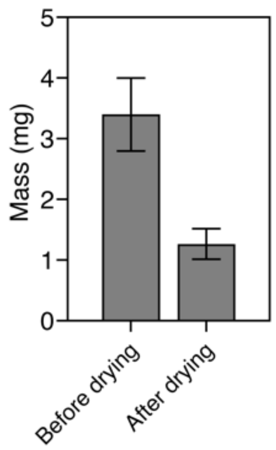


**Figure S15.** Mass of 1-day grown ELM-devices before and after drying. ELM-devices were dried in room temperature for 24 h.

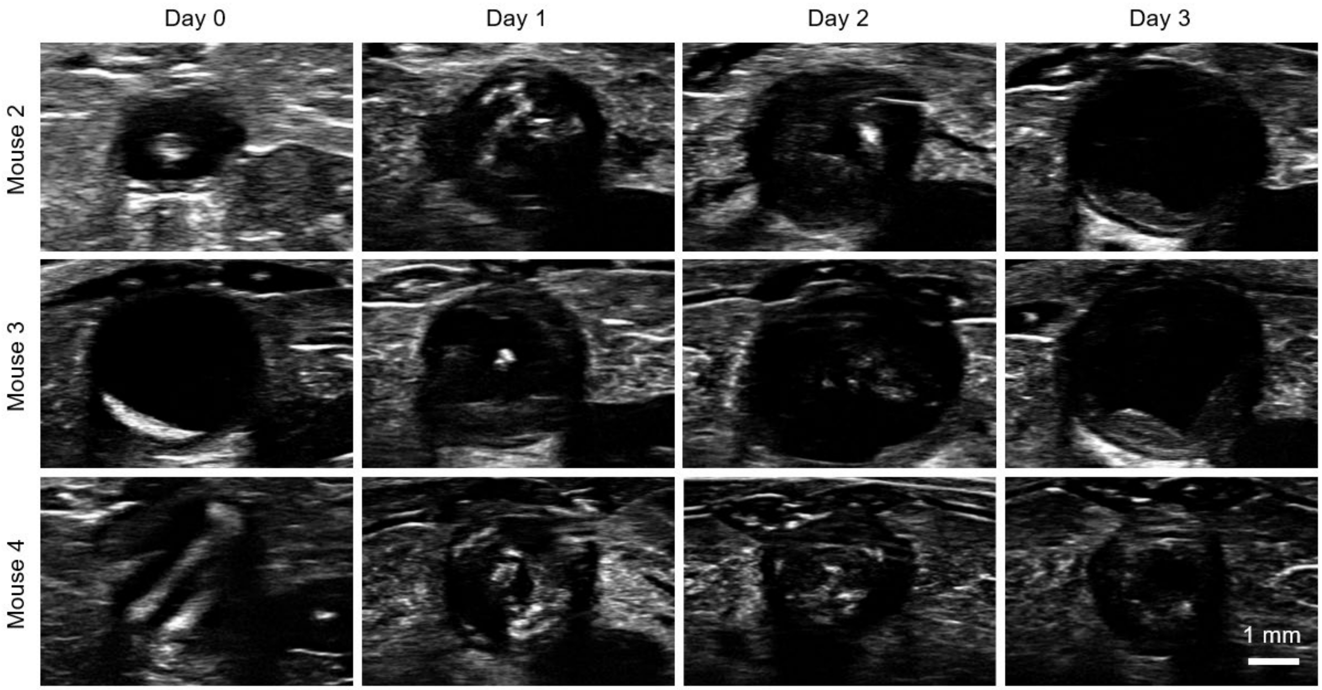


**Figure S16.** Ultrasound images showing device retention in the bladder of mice (Mouse # 2, 3, and 4), each implanted with two ELM-devices. Day 0 images were acquired immediately after implantation, followed by imaging every 24 h for 3 days. All mice retained ELM-devices through day 2. ELM-devices were no longer observed in Mouse 2 and Mouse 3 on day 3, whereas Mouse 4 retained its ELM-devices.

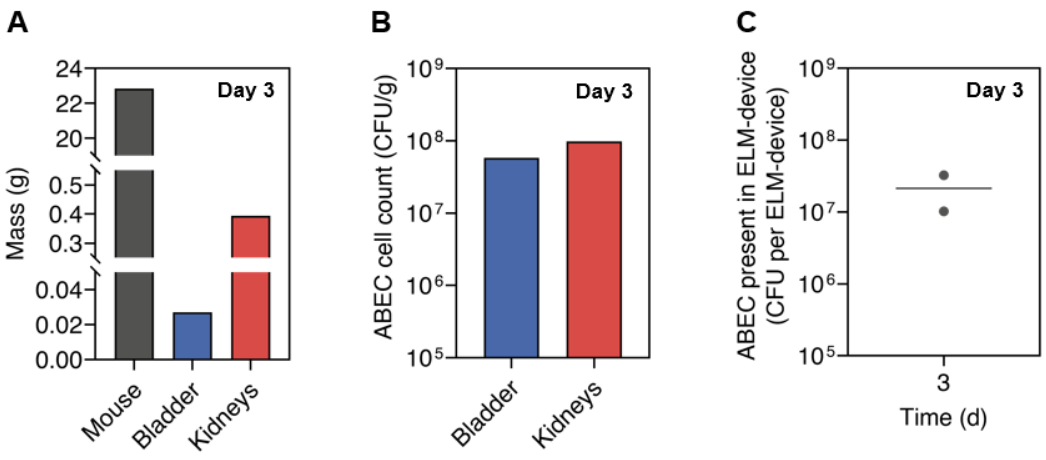


**Figure S17. (A)** Body mass and tissue mass (bladder and kidneys) of a mouse euthanized 3 days post-implantation. **(B)** ABEC counts in the bladder and kidneys of the same mouse. **(C)** ABEC count in ELM-devices recovered from the bladder (*n* = 2). Four ELM-devices were collected from two mice.

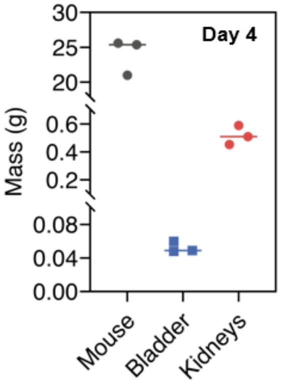


**Figure S18.** Mass of mice and their harvested organs (bladder and kidneys), measured following euthanasia 4 days post-implantation ($n$ = 3).
